# PASTE2: Partial Alignment of Multi-slice Spatially Resolved Transcriptomics Data

**DOI:** 10.1101/2023.01.08.523162

**Authors:** Xinhao Liu, Ron Zeira, Benjamin J. Raphael

## Abstract

Spatially resolved transcriptomics (SRT) technologies measure mRNA expression at thousands of locations in a tissue slice. However, nearly all SRT technologies measure expression in two dimensional slices extracted from a three-dimensional tissue, thus losing information that is shared across multiple slices from the same tissue. Integrating SRT data across multiple slices can help recover this information and improve downstream expression analyses, but multi-slice alignment and integration remains a challenging task. Existing methods for integrating SRT data either do not use spatial information or assume that the morphology of the tissue is largely preserved across slices, an assumption that is often violated due to biological or technical reasons. We introduce PASTE2, a method for *partial* alignment and 3D reconstruction of multi-slice SRT datasets, allowing only partial overlap between aligned slices and/or slice-specific cell types. PASTE2 formulates a novel *partial* Fused Gromov-Wasserstein Optimal Transport problem, which we solve using a conditional gradient algorithm. PASTE2 includes a model selection procedure to estimate the fraction of overlap between slices, and optionally uses information from histological images that accompany some SRT experiments. We show on both simulated and real data that PASTE2 obtains more accurate alignments than existing methods. We further use PASTE2 to reconstruct a 3D map of gene expression in a Drosophila embryo from a 16 slice Stereo-seq dataset. PASTE2 produces accurate alignments of multi-slice datasets from multiple SRT technologies, enabling detailed studies of spatial gene expression across a wide range of biological applications.

**Code availability:** Software is available at https://github.com/raphael-group/paste2

## 1 Introduction

Spatially resolved transcriptomics (SRT) technologies measure mRNA expression simultaneously at thousands of locations within a tissue. These technologies include both sequencing based approaches, such as 10X Genomics Visium [1] and Slide-seq [37, 40], as well as hybridization and florescent approaches such as MERFISH [13] and seqFISH [30]. Nearly all of these technologies measure expression at 2D locations within a thin tissue slice (≈ 10*μm*), and we will use the term spatial transcriptomics (ST) as a generic term to refer to any of these technologies. ST provides spatial context that is missing from single-cell RNA-sequencing (scRNA-seq) measurement of mRNAs from disassociated cells, and has been widely used to study both normal [2, 45] and diseased tissues, such as cancer [23, 39, 42] and Alzheimer’s disease [14]. However, similar to scRNA-seq, ST data suffers from high rates of sparsity. Moreover, recording only the *x, y* coordinates on a 2D tissue slice loses information along the *z* (orthogonal) direction of the 3D tissue, hindering a comprehensive analysis of the whole tissue (Fig. 1).

**Figure 1:**
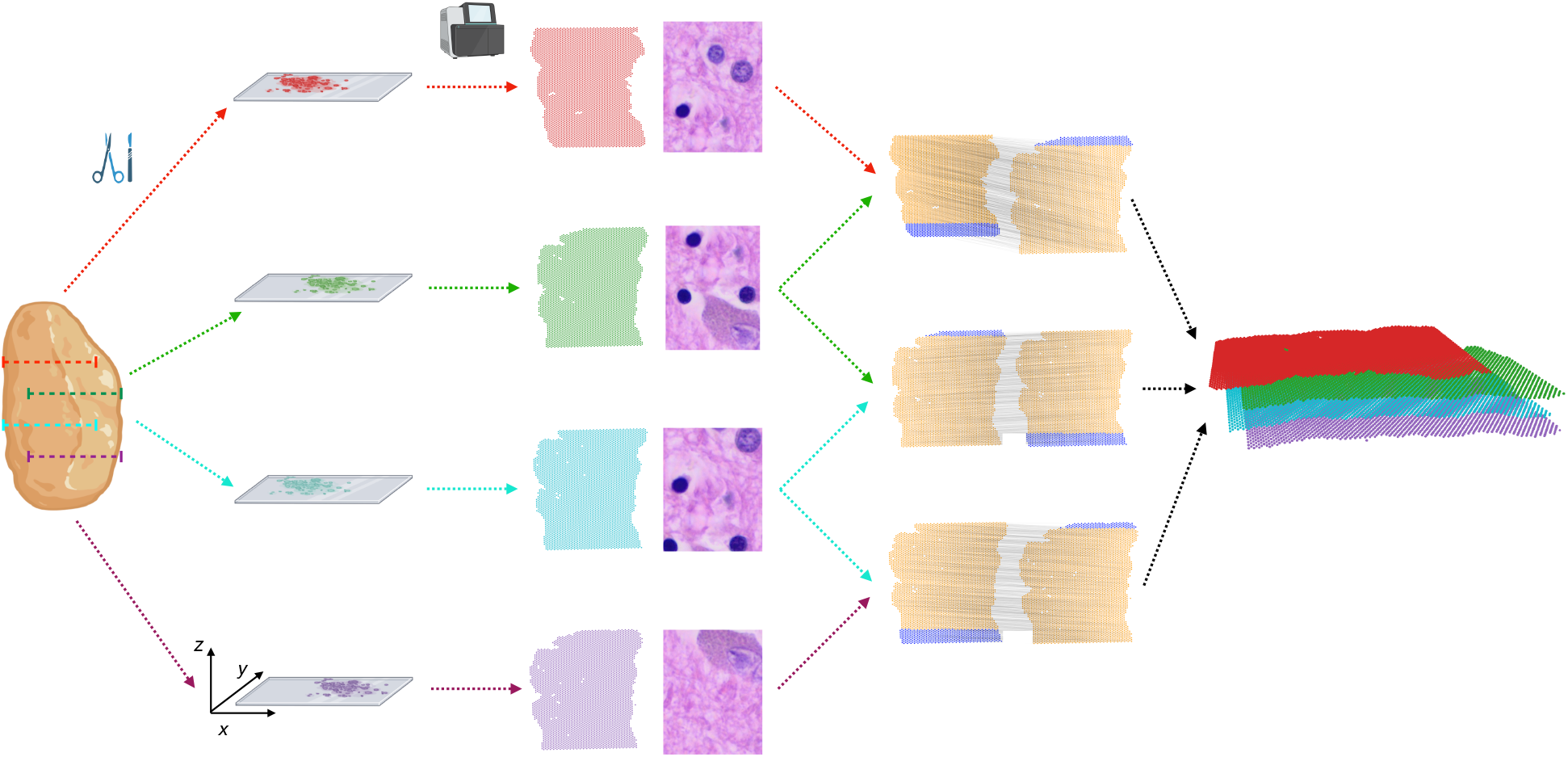
PASTE2 partial alignment of overlapping slices. Four thin slices (red, green, blue, purple) are dissected from the same tissue and placed on an ST array. However, these slices only partially overlap in the *z*-coordinate direction. The inputs to PASTE2 are the four ST slices, including gene expression, spot locations, and optionally, histology images. PASTE2 computes a *partial* alignment of each pair of adjacent slices by selecting subsets of spots from each slice that preserve transcriptional, spatial and image similarity. PASTE2 uses the partial alignment to create a 3D spatial reconstruction of the tissue.

Spatial transcriptomics is often applied to multiple sequential 2D slices from the same tissue (Fig. 1), thus opening the possibility of performing integrative analysis of all slices. Such joint analysis of multiple slices not only helps with the data sparsity problem in individual slices, but also enables innovative downstream tasks such as 3D spatial expression analysis, 3D cell-cell communication, and 3D clustering [29, 48]. However, aligning multiple slices from the same tissue along the orthogonal direction to recover spot-spot correspondence across slices is a challenging task due to morphological differences across slices as well as technical variability in mRNA capture between experiments.

Several approaches have been used for alignment of multiple ST slices. One approach is to apply methods developed for scRNA-seq and multi-omics data integration, such as Seurat [41], SCOT [15, 16], or Pamona [10]. Another approach is to use methods that align an scRNA-seq dataset onto an ST dataset, such as Tangram [5] or RCTD [8]. However, these methods are designed for different alignment tasks and ignore the spatial information within or across slices. Another method, STUtility [4] is designed to align a pair of ST slices, but aligns only the histology images, ignoring both gene expression and spatial information. Morover, this method can only be applied to 10X Genomics Visium data. Another recent method, GPSA [24] integrates multiple ST slices into a common coordinate system, but does not output a mapping between spots that can be used for downstream analysis, and the common coordinate system it produces is different from the 3D coordinates of the tissue.

Another possible solution is to use histological and medical image registration [31] toolkits such as ITK [33] and SimpleITK [3]. However, many image registration methods are supervised and often require manually selected landmarks, creating an extra burden on the user. The spatial alignment problem has also been studied in the context of functional magnetic resonance imaging (fMRI) data registration [7, 26, 27], but these methods are not easily extensible to the spatial genomics setting [24]. Finally, many ST technologies do not have matching histological images.

Recently, PASTE, a method that performs probabilistic alignment of ST slices using both spatial and transcriptional similarities, was introduced [48]. However, PASTE assumes that the slices overlap over the full 2D assayed region, with similar field of view and similar number and proportion of cell types. Essentially, PASTE assumes the two slices are biological/technical replicates of a 2D tissue, an assumption that is often violated in real ST experiments due to technical difficulties in tissue dissection and array placement, or differences in tissue morphology between nearby slices. For example, two slices may only partially overlap along the *z* axis due to different placements of the array on the tissue, and hence only a part of both slices should be aligned (Fig. 1). Furthermore, two slices may have different compositions of cell types, leading to slice-specific cell types and spots that should not be aligned.

Here, we introduce PASTE2, a method to align multiple adjacent ST slices from the same tissue with several substantial improvements over existing methods. First, PASTE2 performs *partial* pairwise alignment, selecting and aligning only a subset of spots. PASTE2 thus addresses the important case where adjacent slices do not fully overlap in space or have different cell type compositions. To solve the partial alignment problem, we introduce the *partial Fused Gromov-Wasserstein (partial-FGW*) *optimal transport* framework. Partial-FGW is the partial extension [11] of the Fused Gromov-Wasserstein optimal transport [43] and allows only a fraction of the total mass to be transported between the two distributions. To the best of our knowledge, PASTE2 is the first to formulate the partial-FGW problem and provide an optimization procedure to solve this problem. Second, PASTE2 includes a model selection procedure to estimate the fraction of overlap between two slices to align, which is in general a very difficult problem. Third, PASTE2 optionally uses the histological images. Some ST technologies, such as the 10X Genomics Visium platform, can also produce a Hematoxylin and Eosin (H&E) stained image of the same tissue slice where gene expression is measured. The information in this image can aid in alignment of slices by identifying spots with similar histology. Finally, we provide a generalized Procrustes analysis [46] method for 3D spatial reconstruction of the tissue from partially aligned 2D slices.

We demonstrate PASTE2’s advantages on both simulated and real ST datasets. We show on simulated data that PASTE2 achieves accurate alignment and outperforms PASTE when slices do not fully overlap. On ST dataset from the human dorsolateral prefrontal cortex (DLPFC) [32], we show that PASTE2 computes more accurate alignments than competing methods, and the use of histological images can further improve the alignment. Finally, we demonstrate PASTE2’s applicability to larger datasets using different SRT technologies by aligning 16 Stereo-Seq slices from a Drosophila embryo [47].

## 2 Methods

A spatial transcriptomics (ST) experiment on a 2D tissue slice yields a pair (*X, Z*), where 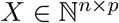 is the gene expression matrix of the tissue slice, and 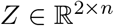 is the spatial location matrix of each spot on the slice, where the *j*-th column ***z**_.j_* is the *x-y* coordinate of spot *j* on the 2D array^1^ used by the ST experiment. Here, *n* is the number of spots on the slice and *p* is the number of genes measured. 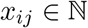 is the transcript count of gene *j* in spot *i*. Each row vector **x**_*i.*_ of *X* is the *expression profile* of spot *i*. Following [48], we encode the spatial location of each spot in a pairwise distance matrix 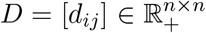, where *d_ij_* is the Euclidean distance between spot *i* and spot *j* on the slice, calculated from the 2D coordinates **z**_.*i*_ and **z**_.*j*_. Thus, we represent an ST slice of *n* spots and *p* genes by a tuple (*X, D*).

### 2.1 Partial pairwise slice alignment problem

Given a pair, (*X, D*) and (*X’, D’*) of ST slices, our goal is to compute a *partial pairwise slice alignment*, i.e. to find a probabilistic spot-spot correspondence between spots in the two slices while accounting for the fact that some spots should not be mapped (Fig. 1). The probabilistic mapping is a matrix 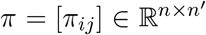 between the *n* spots in one slice and *n’* spots in the other slice, where *π_ij_* describes the probability (or relative fraction) that a spot *i* in the first slice is aligned to a spot *j* in the second slice.

We begin by describing the solution given in [48] to the pairwise slice alignment problem, implemented in the PASTE algorithm. PASTE uses a formulation based on optimal transport to compute the mapping *π*. Specifically, given probability distributions *g* and *g’* over the spots in slice *X* and *X’*, respectively, PASTE finds the map *π* (also known as the transport matrix) that minimizes the following transport cost:

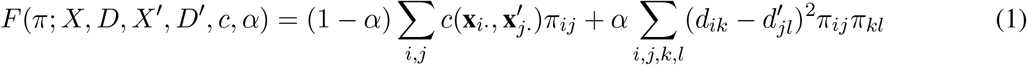

subject to the regularity constraint that *π* has to be a probabilistic coupling between *g* and *g’:*

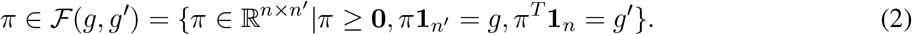

Here, 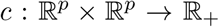 is an *expression cost function* that gives a non-negative dissimilarity score between the expression profiles of two spots over the same genes. **1**_*n*_ is an all-one vector of length *n*. Typically, *g* and *g’* are chosen to be uniform distributions over spots in each slice, although other distributions can be used [48].

The PASTE objective function *F* is composed of an expression similarity term (first summand) and a spatial similarity term (second summand) weighted by a parameter *α*. The first term, also called the Wasserstein distance in the OT literature [35], represents the cost of moving one unit of probability mass from each spot *i* to each spot *j*, with the cost being the gene expression dissimilarity between spots. The second term, also called the Gromov-Wasserstein distance [34, 36], approximately preserves the intra-slice spatial distances between spots. Together, the convex combination of the two terms in *F* is known as the Fused Gromov-Wasserstein (FGW) optimal transport objective [43].

The regularity condition (2) forces a rigid structure on *π* such that all spots from both slices must be aligned. However, such constraints may not be appropriate for ST slices with considerable differences in field of view or cell type composition due to both biological variation across tissue sections as well as differences caused by the manual nature of tissue dissection. Therefore, spots containing cell types or tissue regions that are unique to only one slice will be forced to be mapped to somehow arbitrary spots on the other slice.

Thus, in PASTE2, we propose to solve the *partial pairwise slice alignment problem* by minimizing the same objective function as PASTE (Equation (1)), but with a different set of constraints that allow for unmapped spots. Specifically, given a parameter *s* ∈ [0,1] describing the fraction of mass to transport between *g* and *g’*, we define a set 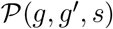 of *s-partial* couplings between distributions *g* and *g’* as

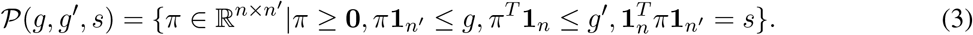

The parameter *s* ∈ [0,1] is interpreted as the overlap percentage between the two slices to align. The constraint 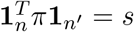 ensures that only the fraction of s probability mass is transported. Equivalently, if 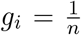 is a point mass for each spot, then roughly s fraction of the spots in each slice are aligned. The feasibility constraints π ≥ 0, ∑_*j*_ *π_ij_* ≤ *g_i_* for all spots *i* in the first slice, and 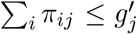 for all spots *j* in the second slice make sure that each spot only transport probability mass that it already has according to *g* and *g’*, hence ensures *π* to be a valid transport plan. In PASTE2 we require that the map *π* belong to 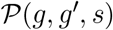, thus replacing the set 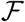 (defined in (2)) by the set 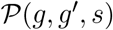 (defined in (3)). In analogy to sequence alignment, PASTE calculates a global alignment, while PASTE2 calculates a local alignment.

The general concept of partial optimal transport [9] extends optimal transport theory to allow the transportation of only a specified fraction of mass between distributions. Here, we adapt the idea of partial optimal transport to the Fused Gromov-Wasserstein objective, hence the PASTE2 optimization problem is a novel partial Fused Gromov-Wasserstein (partial-FGW) optimal transport problem. While there are existing solutions to the partial Wasserstein and partial Gromov-Wasserstein problem, to the best of our knowledge, PASTE2 is the first to state and formulate the partial-Fused Gromov-Wasserstein problem and provide an optimization procedure to sovle this problem.

PASTE2 has two parameters: *α*, the balance between the gene expression dissimilarity and the spatial dissimilarity, and the overlap percentage parameter *s* indicating the fraction of mass to transport. Unless otherwise specified, we set *α* = 0.1 following [48]. We choose the value of *s* using a model selection procedure described in Supplement §1. The choice of *c*, the expression dissimilarity function is described in Supplement §2.

### 2.2 An iterative conditional gradient algorithm for optimization

We derive an optimization algorithm to minimize the objective (1) subject to the constraint (3). This problem is a large scale (each slice contains thousands of spots) non-convex quadratic program with a convex and compact feasible region. Our algorithm is based on the Frank-Wolfe optimization algorithm [20], also known as the conditional gradient [28] algorithm. This algorithm has been widely adopted in the optimal transport community [11, 17, 19, 43] to compute transport plans because of its ability to handle large-scale quadratic programs [22]. The optimization problem in PASTE2 is thus particularly suitable for the conditional gradient algorithm.

The conditional gradient algorithm is an iterative first-order algorithm for constrained optimization. To fit in the conditional gradient scheme, we first write (1) in matrix form, following [36]

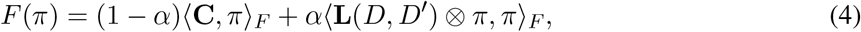

where 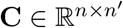 encodes the gene expression dissimilarity 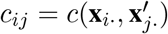 between each spot *i* in the first slice and each spot *j* in the second slice, and 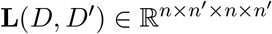 is a 4-dimensional tensor defined by 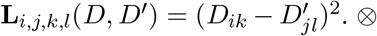 is the tensor-matrix multiplication operator, *i.e*. **L** ⊗ *π* is an *n* × *n’* matrix whose (*i*, *j*)-th element is (∑_*k,l*_ **L**_*i,j,k,l*_ · *π_k,l_*). 〈·, ·〉) denotes the Frobenius dot product of matrices.

In each iteration, the algorithm moves in the direction that minimizes a linear approximation of the objective function while remaining in the feasible region. The mathematical details of the derivation of each step, as well as the pseudocode, is provided in Supplement §3.

### 2.3 Using histological image data in alignment

We further extend the PASTE2 partial-FGW framework to incorporate image information. Specifically, we replace the gene expression dissimilarity matrix 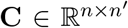 in Equation (4) by a sum of two *n* × *n’* dissimilarity matrices 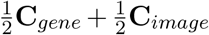 where **C**_*gene*_ is the gene expression dissimilarity matrix as defined above and **C**_*image*_ encodes the dissimilarity between the image information at each spot. Thus, we seek a map π that minimizes the following objective function

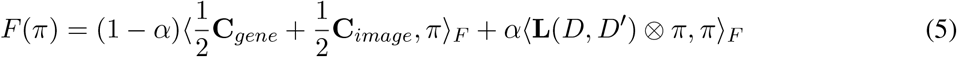

Note that to avoid an extra parameter we give equal weight 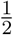 to both gene expression and image information, although substituting other weights is straightforward. Also, since **C**_*gene*_ and **C**_*image*_ may not be on the same scale, we scale **C**_*image*_ such that the maximum entry of **C**_*image*_ equals the maximum entry of **C**_*gene*_. In our implementation, we define [**C**_*image*_]_*ij*_ to be the Euclidean distance between the mean RGB values of the spots. See Supplement §4 for further details.

### 2.4 3D reconstruction based on the partial alignment matrix

Given a series of consecutive, (partially) overlapping slices from the same tissue, we aim to reconstruct the spatial expression of the tissue in 3D by transforming PASTE2 partial pairwise alignments into a common coordinate system. Specifically, given a series of consecutive slices we first find partial alignments between adjacent slices by solving the partial pairwise slice alignment problem as above. To project all slices onto a common coordinate system, we extend the generalized weighted Procrustes analysis [25, 46] approach in [48] to sequentially project each pair of adjacent slices. While [48] projects a pair of slices onto the same coordinate system by centering both slices followed by calculating a rotation matrix, we derive the centering step for each slice separately to address the case where the alignment matrix *π* is partial and the aligned regions of the two slices have unique barycenters. The details of the projection are in Supplement §5.

## 3 Results

### 3.1 Evaluation on simulated ST data

We first compared PASTE2 and PASTE on a simulated ST dataset based on a human dorsolateral prefrontal cortex (DLPFC) tissue slice from [32]. Specifically, we extracted two partially overlapping subslices from a single DLPFC slice (sample 151674, corresponding to Slice 3B in §3.2) with varying overlap percentages 90%, 70%, 50%, 30% (Fig. 2a). To perturb the gene expression, we resample the gene expression profile of each spot in one of the subslices by sampling from a multinomial distribution with added pseudocount *δ*, which controls the noise level (Supplement §6). We vary the pseudocount *δ* in the range^2^ from 0.1 to 2 with an increment of 0.1. In total, we generated 4 × 30 = 120 pairs of subslices with different overlap percentages and noise levels *δ*. For each pair of subslices, we ran PASTE2 with *α* = 0,0.1,1 and using the ground truth value for the overlap fraction *s* (we evaluated model selection separately in Supplement §9), as well as full PASTE with default parameters (*α* = 0.1). We evaluated the alignment using *Label Transfer Adjusted Rand Index (LTARI*). Given a labeling of cell type/spatial region of spots, LTARI measures how well the alignment preserves the label between the aligned spots. LTARI first defines a new spot labeling for the second slice by assigning to each aligned spot the label of the most likely corresponding spot in the first slice, then calculates the ARI between the induced spot labeling of the second slice and the ground truth labeling (Supplement §7). We used the manual cortical layer annotation from [32] as ground truth spot labeling (Fig. 2a).

**Figure 2:**
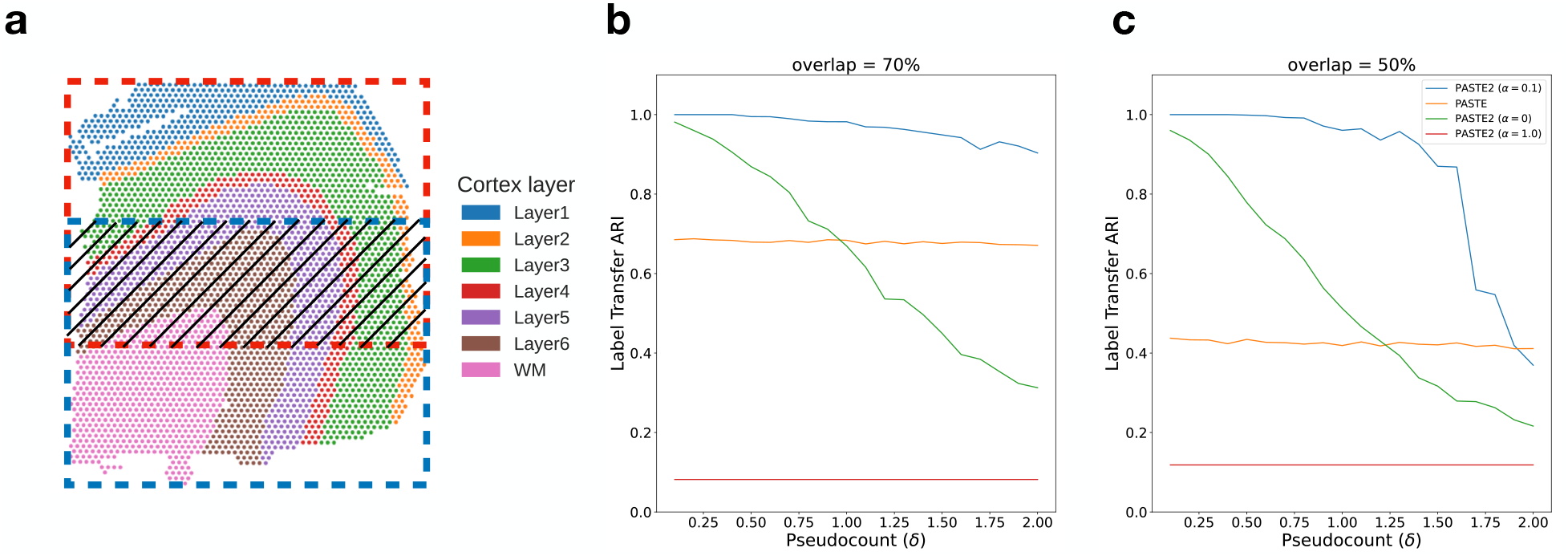
Comparison of PASTE2 and PASTE on simulated partially overlapping subslices. **a**, DLPFC slice 151674 with spots colored according to the manual annotations of cortical layers from [32]. Red box and blue box indicate two partially overlapping subslices, with central overlapping region containing some fraction of spots from each slice. **b**, Label Transfer ARI of the alignments produced by PASTE2 with *α* = 0 (gene expression information only), PASTE2 *α* =1 (spatial information only), PASTE2 *α* = 0.1 (both), and PASTE (full alignment, *α* = 0.1) as a function of the pseudocount (*δ*) for overlap percentage70%. **c**, Label Transfer ARI for overlap percentage 50%.

We found that PASTE2 with the default parameter setting of *α* = 0.1, which uses both gene expression information and spatial information, outperforms PASTE across most values of the added noise *δ* for every overlap percentage (Fig. 2b, Fig. S6). Specifically, for all four overlap percentages, PASTE2 (*α* = 0.1) achieves the highest LTARI when *δ* < 2.0, and achieves almost perfect LTARI when *δ* is small. Note that PASTE obtains constant accuracy because it aligns overlapping regions well but non-overlapping regions arbitrarily. The gap in accuracy between PASTE2 and PASTE is larger when the overlap is smaller. This indicates that PASTE, which finds an alignment between all pairs of spots, is not suitable for the partial alignment task. In contrast, PASTE2 has high accuracy in partial alignment across a wide range of overlap percentages and gene expression noises. Moreover, PASTE2 achieves near perfect LTARI when the added pseudocount is in the range of variability in read counts (≈ 0.1 – 0.2) observed in real data [48].

To investigate the effect of the misspecification of the value of the overlap percentage parameter *s* on the result of PASTE2, we ran PASTE2 with *s* ranging from 0.1 to 1, with a step size of 0.1, on a simulated pair where the ground truth overlap percentage is 50% and the added pseudocount is 0.1. We found that PASTE2 aligns correctly when *s* is lower than the ground truth, while the performance degrades for larger values of *s* (Fig. S7). This is expected because with an overestimation of the overlap percentage, the PASTE2 alignment becomes more similar to the PASTE alignment which includes all the spots. Thus, in selecting a value for *s*, it is preferable to use a model selection procedure that slightly underestimates *s* rather than overestimate s. We propose a heuristic for selecting s in Supplement §1.

Finally, we emphasize the importance of using both gene expression and spatial information in computing accurate partial alignments. PASTE2 (*α* = 1.0) has consistently low LTARI, indicating that using spatial coordinates alone cannot recover alignment across slices. The performance of PASTE2 (*α* = 0), which only uses gene expression information of each spot for alignment, drops more quickly than PASTE2 (*α* = 0.1) with increasing pseudocount *δ* indicating more noise in gene expression. Using both gene expression and spatial information, PASTE2 is able to accurately align two partially overlapping ST slices. The effect of different intermediate values of *α* on the alignment performance is thoroughly discussed in [48].

### 3.2 Human dorsolateral prefrontal cortex (DLPFC) slices

We next compared PASTE2 to PASTE [48] and two other transcriptomics alignment methods – Pamona [10] and Tangram [5] – on the full human dorsolateral prefrontal cortex (DLPFC) dataset containing 10X Genomics Visium ST data from three individuals (labeled sample 1, 2, 3) with four slices (labeled slice A, B, C, D) per individual [32]. For each individual, slices A and B and slices C and D are 10 *μm* apart. However, slices B and C are further apart at a distance of 300 *μm*. Hence, slice pairs AB and CD are more similar to each other than slice pair BC. Note that Pamona [10], a manifold alignment algorithm for multi-omics datasets, is also based on partial optimal transport, while Tangram [5] is a deep-learning based method that aligns scRNA-seq data onto ST data. We create partial ST alignment problems by generating two partially overlapping DLPFC datasets as follows. For each individual, we extracted the left portions of slice A and C, and the right portions of slice B and D such that the extracted pairs AB, BC, and CD have ≈ 70% overlap in area (Fig. 3a). We also created another set of partially overlapping dataset using horizontal slices (Fig. S8a). We ran each of the methods as described in Supplement §8. We evaluate the accuracy of each method by computing the Label Transfer ARI (LTARI) as previously described (Section 3.1).

**Figure 3:**
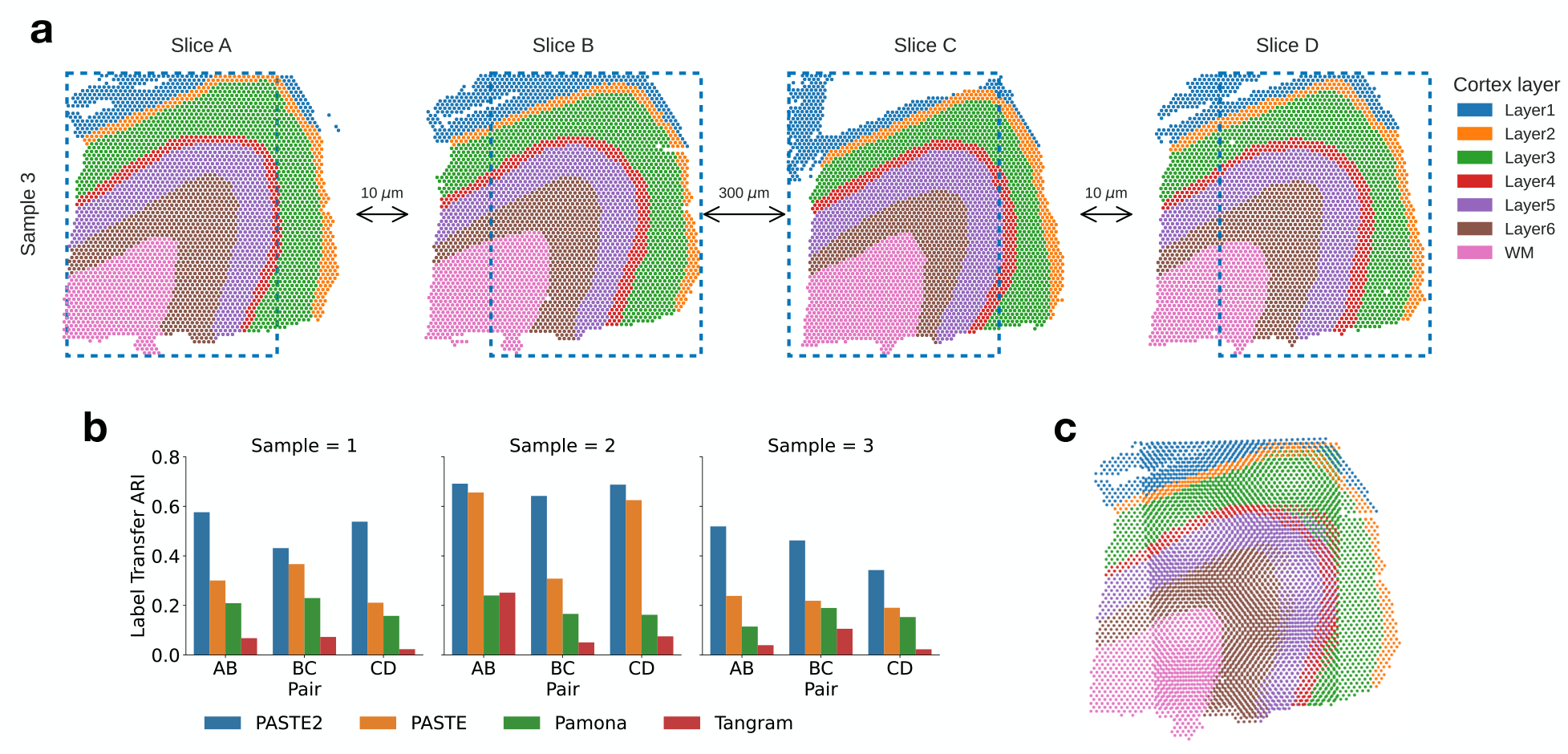
Comparison of alignment methods on partially overlapping DLPFC slices. **a**, Vertical subslices were obtained by cropping subslices (blue dotted boxes) from four adjacent slices from DLPFC sample 3, with indicated distances between adjacent slices. Each pair of adjacent subslices overlaps in 70% of their areas. **b**, LTARI of pairwise alignments computed by PASTE2, PASTE, Pamona, and Tangram for each pair of adjacent vertical subslices from three samples. **c**, Optimal projection of vertical subslices from slice AB of sample 3 onto the same 2D coordinate system using the PASTE2 partial alignment.

We find that PASTE2 achieves the highest LTARI on all adjacent subslices of all individuals, for both vertical partial slices and horizontal partial slices, with the exception of one pair (Fig. 3b, Fig. S8b). About 70% - 75% spots from each subslice is aligned in each pair indicating the parameter *s* corresponds well with slice overlap. For most pairs, PASTE2 has more than twice the LTARI than all other methods, demonstrating PASTE2’s ability to identify the overlap region of the two ST slices and align the overlap region reliably. On one pair Pamona has slightly higher LTARI than PASTE2 (Fig. S8b), but all methods have very low LTARI (< 0.1), suggesting that this pair has low spatial coherence. PASTE is the second-highest performing method on most pairs, indicating that even though PASTE does not model partially overlapping slices, it is still more suitable for aligning spatial transcriptomics data than methods designed for different purposes. While Pamona is designed to align datasets with both shared and dataset-specific cells [10] – the analog of the partial pairwise slice alignment problem for single cell datasets – Pamona does not model spatial constraints, perhaps explaining its lower performance. Tangram assumes the single-cell gene expression dataset and the spatial dataset come from the same anatomical region [5], hence the partial slice alignment task violates the Tangram assumption, leading to a low alignment accuracy.

For a more intuitive demonstration of PASTE2’s advantage and accuracy, we projected the vertical subslices of sample 3 pair AB onto the same coordinate system, computed as described in §2.4 based on the alignment matrix computed by PASTE2 (Fig. 3c), as well as the optimal projection of the same pair based on the alignment computed by PASTE (Fig. S9a). Qualitatively, the projection of PASTE2 correctly stacks the overlap area of the two slices, with spots from the same cortical layer stacking on top of each other, while PASTE fails to find the corresponding layers in the two slices. Additionally, PASTE2 correctly identifies and aligns the overlap area of all four partial slices of an individual while leaving the rest unaligned, leading to a visually correct 3D reconstruction of the tissue from partial slices (Fig. S9b).

We also ran STUtility [4], a method to align H&E stained images that are generated as part of the 10X Genomics Visium ST workflow. STUtility outputs new coordinates of the aligned slices and does not produce a mapping between pairs of spots; thus, we visualized the alignment results by plotting each pair of partial subslices according to the new coordinates output by STUtility (Fig. S10). The image masking function utilized by STUtility failed for the partial slices of sample 3, so we only visualized the results for sample 1 and 2. STUtility correctly identifies that each pair of input slices are partially overlapping, but it does not align the correct overlapping region, and the output alignment seems quite arbitrary. These results might be because STUtlitiy aligns images by identifying and finding correspondences between edges of the two input tissues, but when two tissues are partially overlapping, the edges do not provide information about spot correspondences. On the other hand, PASTE2 correctly aligns the overlapping region (Fig. 3c). This demonstrates that using transcriptomic similarity, spatial similarity, and image information yields better partial alignments than H&E images alone.

Finally, we compared PASTE2’s running time with other methods on the vertical subslices of sample 3. PASTE2 finished in under 10 minutes for all subslice pairs on a Macbook Pro with 2.4GHz Intel Core i5 CPU, with most of the running time spent on the GLM-PCA subroutine (Fig. S11). The conditional gradient optimizer in PASTE2 runs in less than half of the time of Pamona and Tangram, and only runs slightly slower than PASTE. We also used the DLPFC datasets to evaluate the accuracy of PASTE2’s model selection procedure for estimating the overlap percentage s, and found that PASTE2 correctly estimates the overlap percentage in many scenarios (Supplement §9).

### 3.3 Incorporating histology information improves alignment

We compared PASTE2’s alignment performance when using both gene expression and histological image (Equation 5) versus using only gene expression data (Equation 4). Note that spatial information is included in both analyses. We ran the two modes of PASTE2 on pairs of horizontal and vertical subslices from DLPFC sample 3. We found that using the histological image substantially improved the alignment performance for pair CD (Fig. 4ab), increasing the LTARI from 0.34 to 0.46. Examining the alignment obtained on this pair using only gene expression information (Fig. 4c) to the alignment obtained with both gene expression and the histology image (Fig. 4d), we observe that the alignment obtained using the images is more spatially contiguous. In particular, there is a curve of unaligned spots (blue spots in Figure 4c) in subslice D inside the yellow region. This curve corresponds to spots that are manually annotated as Layer 6 of the DLPFC [32]. Interestingly, the spots in this layer have lower total UMI counts than other layers: the mean total UMI counts for spots from Layer 6 is 2915 compared to total UMI counts ≈ 4500 in the other layers. This suggests that the gene expression signal is weaker in these spots. In contrast, the PASTE2 alignment obtained using both gene expression and image information (Figure 4d) does not have the same curve of unaligned spots, demonstrating the advantages of using the histological image for spots with a weak gene expression signal.

**Figure 4:**
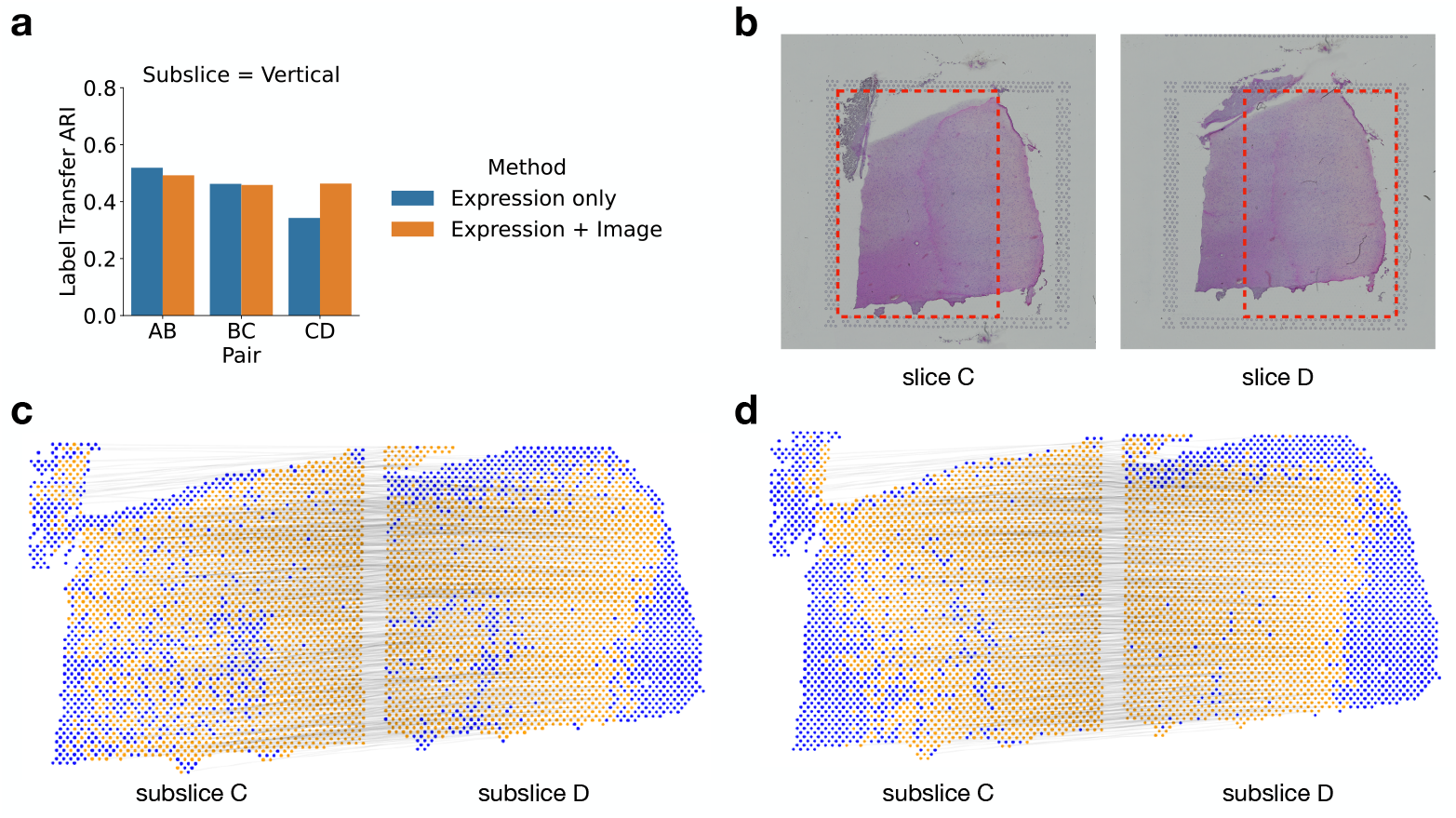
Evaluating the benefit of using histological image information in PASTE2 alignment. **a**, The Label Transfer ARI (LTARI) of PASTE2 partial alignments of pairs of vertical subslices extracted from DLPFC sample 3 using only gene expression (blue) and using both gene expression and image information (orange). **b**, Histological images of sample 3 slice C and slice D. The red boxes bound the vertical subslices extracted for partial alignment. The right part of subslice C should be aligned to the left part of subslice D. **c**, Visualization of PASTE2 alignment of the subslice pair CD using gene expression and spatial information. Yellow spots are aligned by PASTE2, while blue spots are unaligned. Thin black lines connect pairs of spots that are aligned by PASTE2 with high weight. **d**, Visualization of PASTE2 alignment of the same subslice pair when gene expression, histological image, and spatial information are all used.

In the horizontal slices, we see cases where using the image information reduces the alignment performance. For example, the LTARI drops from 0.56 (expression) to 0.50 (expression and image) for horizontal subslices of pair AB (Fig. S14a,b). The aligned part of subslice B in Fig. S14d shows that many spots are left unaligned in the actual overlap region, and there is a clear stripe of unaligned blue spots towards the left part of the subslice. Looking at the H&E image of subslice B in Fig. S14b, we see a clear dark stain on the left of the subslice that is missing from the image of subslice A, at exactly the same location of the unaligned stripe. This indicates that the stain on the H&E image is the cause for the worse alignment performance.

For the other pairs, the LTARI for PASTE2 alignments with and without images is approximately the same. This is not too surprising since the H&E images of DLPFC slices do not display strong heterogeneity across different layers (Fig. 4b, Fig. S14b). However, the fact that utilizing image information corrects the alignment of low UMI spots demonstrates the potential for histological images to guide PASTE2 alignment. The image information can help overcome the sparsity of gene expression, and when the histological images have greater variation across spots, using the images should further improve the alignment quality by complementing the gene expression signal.

### 3.4 Spatial transcriptomics of Drosophila embryo

We applied PASTE2 to analyze a Stereo-seq dataset from a Drosophila embryo [12]. Stereo-seq is a new SRT technology with ≈ 500nm resolution, two orders of magnitude smaller than the 10X Visium platform, but with lower UMIs per spot. [47] applied Stereo-seq to two late-stage Drosophila embryos 14-16 hours and 16-18 hours after egg laying (labeled E14-16 and E16-18) and three stages of larvae (labeled L1-L3). Each slice has ≈ 1000 spots with median UMI per spot of ≈ 2000, compared to ≈ 4000 spots and ≈ 5000 median UMI per spot in the 10X Visium DLPFC dataset. In the published analysis, the cell type of each spot was derived by unsupervised clustering of gene expression followed by annotation based on marker genes. The publication used PASTE to align all slices from the same stage and obtain a 3D map of spatial expression of each stage. However, slices from the same stage vary in size and cell type compositions and do not fully overlap in space. For example, inspection of annotated cell types shows that adjacent slices from the E14-16 sample do not fully overlap (Fig. S15). Therefore, it is appropriate to use PASTE2 to realign the adjacent slices respecting the different composition of cell types across slices, and to obtain a more accurate 3D reconstruction of the Drosophila embryo.

We applied PASTE2 to compute a partial alignment for each pair of 16 adjacent slices from the E14-16 sample, estimating the overlap percentage using the PASTE2 model-selection heuristic (Supplement §1). Slices 7 and 8 have clear differences in the composition of cell types annotated by [47], with the carcass cells showing the largest difference in proportion (Fig. 5a). PASTE2 addresses this imbalance by aligning a similar proportion of carcass cells across slices, leaving the excess cells in slice 8 unaligned. The spots from the two slices included in the PASTE2 partial alignment show similar spatial organization (Figure 5b) and cell type composition (Fig. S16b). For example, the proportions of carcass cells in slices 7 and 8 differ by 10% before alignment (Fig. S16a), but after alignment the difference is less than 3% (Fig. S16b). The differences in proportions shrinks for salivary gland cells as well, indicating PASTE2 correctly identifies and aligns the overlapping parts. PASTE2 optimal projection of the two slices to the same coordinate system puts slice 8 slightly higher in *y* coordinates than slice 7, consistent with the observation that slice 8 has unaligned carcass cells at the top (Figure 5c). The LTARI obtained by PASTE2 for this pair is 0.49, compared to a LTARI of 0.39 for PASTE, again showing the advantages of partial alignment. Examination of pair of adjacent slices 14 and 15 shows a similar advantage of partial alignment. Slice 15 has a stripe of carcass cells that is absent in slice 14 (Fig. S17a). PASTE2 leaves the stripe unaligned across slices (Fig. S17bc), increasing the LTARI from 0.29 for PASTE to 0.52 for PASTE2. Since PASTE computes an alignment for all spots, the extra carcass cells in slice 15 are mapped somewhere on slice 14, creating false correspondences between spots (Fig. S18).

**Figure 5:**
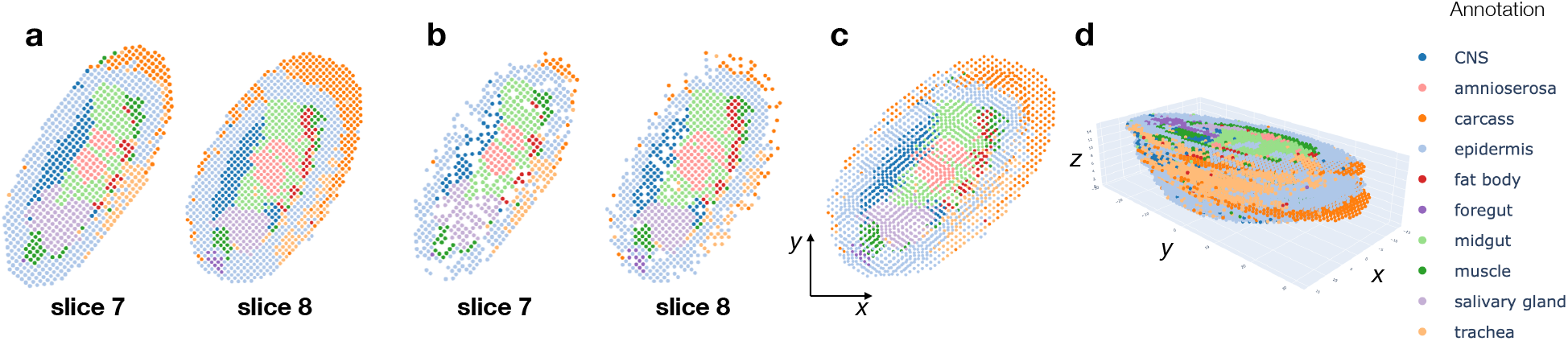
PASTE2 alignment of Stereo-seq data from E14-16 Drosophila embryo from [47]. **a**, Stereo-seq slices 7 and 8 with spots labeled by cell types annotated in [47]. **b**, Spots from slices 7 and 8 that are included in the partial alignment computed by PASTE2. Spots selected by PASTE2 have similar proportions of cell types and spatial locations. **c**, Optimal projection of slices 7 and slice 8 onto the same 2D coordinate system using the PASTE2 partial alignment. **d**, PASTE2 3D reconstruction using all 16 slices from the Drosophila embryo.

We compared the LTARI of the PASTE2 alignment with the LTARI scores of PASTE, Pamona, and Tangram on every pair of adjacent slices. PASTE2 achieves the highest LTARI for most pairs, with the largest gain in pairs where the two slices have different compositions of cell types, such as slice 14 and 15 (Fig. S19). Pairs where PASTE2 does not obtain the highest LTARI, such as slice 2 and 3, have relative similar sizes and cell types, and PASTE2 still achieves comparable LTARI with the highest performing method. This indicates that PASTE2 not only aligns partially overlapping slices correctly, but also performs well on pairs of similar slices.

We used PASTE2 to generate a 3D reconstruction of all 16 slices of the E14-16 Drosophila embryo, where adjacent slices have on average 70% of overlapping spots (Fig. 5d). The PASTE2 3D reconstruction will be useful for refining the analyses presented in [47] who demonstrated that the PASTE-generated 3D expression helped detect functional subregions and uncover the dynamics of cell state changes and tissue-specific gene regulation.

## 4 Discussion

We present PASTE2, a method to perform pairwise alignment and 3D reconstruction of multi-slice spatial transcriptomics data. PASTE2 addresses the important situation where slices partially overlap in space or have different cell type compositions, which is the case for most real datasets. We formulate the ST partial pairwise alignment problem using a partial Fused Gromov-Wasserstein optimal transport framework and derive an optimization algorithm to solve this problem. We further design a model selection procedure to determine the overlap between slices, and extend the framework to incorporate both gene expression and imaging information.

We found that PASTE2 outperforms multiple other methods for alignment of spatial transcriptomics or single cell data including PASTE, Pamona, Tangram, and STUtility. We show that PASTE2’s use of histology images can further improve alignments, although the results are variable depending on the quality of the images. We expect that PASTE2 will achieve much higher accuracy incorporating image information in datasets where histological images display stronger signal across spots – in preliminary results on unpublished cancer datasets with high-quality H&E images we observed even larger gains. Finally, we demonstrate PASTE2’s capabilities on larger datasets from another SRT technology by generating a 3D reconstruction of a Drosophila embryo from 16 slices of Stereo-seq data.

There are multiple directions for future work. First, is to extend the partial alignment framework to integrate multiple slices into a single consensus slice to address the data sparsity issue by pooling counts from corresponding spots [48]. Second, one could stitch together multiple partially overlapping slices into a larger 2D slice. This stitching would be helpful in cases where adjacent tissue slices are close in the z-coordinate which is often the case with thin tissue slices (≈ 10*μm*). In addition, one could incorporate additional spatial regularization terms to enforce more contiguous overlapping regions. Third, it would be interesting to apply PASTE2 to integrated spatial transcriptomics and imaging data from other platforms such as Slide-seq [37, 40], or combined Stero-seq and imaging data which [47] noted as a future technology development. Finally, it would be interesting to examine the effectiveness of other optimal transport frameworks such as unbalanced OT [38] that impose soft constraints rather than hard constraints on partial alignments.

We anticipate that PASTE2 will be a useful tool for integrating transcriptomic information across multi-slice ST datasets and for building 3D tissue atlas across both normal and diseased tissues, such as in the Human Tumor Atlas Network and related projects.

## Footnotes

1 We refer here to array based technologies, but the formulation is the same for other technologies.

2 With the typical sequence coverage and data sparsity in ST data, *δ* > 2.0 (adding > 2 counts to each transcript) is a strong perturbation of the data that essentially destroys the signal present in the original data.

## Acknowledgements

This work is supported by grants U24CA211000 and U24CA248453 from the US National Cancer Institute (NCI).

## Notes

### Competing Interest Statement

The authors have declared no competing interest.

