## Supplementary material for "PASTE2: Partial Alignment of Multi-slice Spatially Resolved Transcriptomics Data"

### SUPPLEMENT

#### 1 Estimation of the overlap percentage

PASTE2 requires a user-specified parameter $s$, the amount of mass to transport, which is interpreted as the percentage of overlap between the two slices to align. However, this parameter is not available in general. In this section, we introduce a heuristic based on the region aligned by PASTE2 to estimate $s$, the true overlap percentage.

Because the second term in the PASTE2 objective prefers symmetrical alignment, we found that if we overestimate $s$ and visualize the regions in the two slices aligned by PASTE2, the two regions will generally be contiguous in each slice, with the true overlap region being a subset of the aligned region. However, if we underestimate $s$, the aligned regions will not be contiguous, where the true overlap region will have random spots left out unaligned. In other words, if we overestimate $s$ and visualize the convex hull of the aligned region, the convex hull will contain mostly contiguous aligned spots, but if we underestimate $s$ and visualize the convex hull of the aligned region, the convex hull will have unaligned spots randomly spread across the region. To quantify the contiguity of aligned spots, we define an *edge inconsistency score* that measures the spatial coherence of a graph with nodes colored by two clusters. Specifically, let $G = (V, E)$ be a graph and let $L = [l(i)]$ be a labeling of nodes where $l(i) \in \{1, 2\}$ is the cluster label of node $i$. Let $E'$ be the subset of edges where the labelling of the nodes at the two ends are different, *i.e.* $E'$ is the cut of the graph. We define the edge inconsistency score as $H(G, L) = \frac{|E'|}{|E|}$, which is the percentage of edges that are in the cut. A high inconsistency score means most of the edges are in the cut, indicating the labeling of the nodes has low spatial coherence, while a low inconsistency score means the two classes of nodes are mostly contiguous in graph.

Given two slices $(X, D)$ and $(X', D')$, we run PASTE2 with $s$ decreasing from 1 to 0.05 with a step size of 0.05. For each $s$, we calculate the edge inconsistency score of the convex hull of the regions that PASTE2 selects for alignment in the two slices, with each spot labeled aligned or unaligned. Ideally, the edge inconsistency score should remain low when input $s$ is higher than the true overlap percentage $s^*$, and increases when $s$ drops below $s^*$, peaking at exactly $\frac{s^*}{2}$. Therefore, we find $s'_1, s'_2$ which respectively achieves the highest edge inconsistency score in the two slices, and estimate the true overlap percentage as $\widehat{s} = 2\min\{s'_1, s'_2\}$.

#### 2 The gene expression dissimilarity function

The PASTE2 objective needs an expression cost function $c : \mathbb{R}^p \times \mathbb{R}^p \to \mathbb{R}_+$ that measures the dissimilarity level of two gene expression profiles that are potentially on the order of tens of thousands in dimension. Instead of using Kullback–Leibler divergence or Euclidean distance between two expression vectors as PASTE does, PASTE2 computes a dissimilarity cost between two high-dimensional expression vectors as follows. PASTE2 first selects the top 2000 genes with the highest UMI counts across both slices. Then, it uses GLM-PCA [44], a generalization of principle component analysis to exponential family likelihoods, to reduce the dimension of the expression vector at each spot from 2000 to 50. The dissimilarity between two 50-dimensional vectors will then be calculated using standard Euclidean distance. GLM-PCA is designed to operate on raw UMI counts based on a multinomial generative model for expression vectors, avoiding the potential pitfalls of common practices such as normalization and log transformation [44, 21], hence particularly suitable for dimensionality reduction in spatial transcriptomics given its nature of sparsity and high technical variations across spots.

#### 3 A conditional gradient algorithm for partial-FGW optimal trasport

As in the classical conditional gradient procedure, we initialize $\pi^{(0)}$ randomly, then for each iteration $k$, PASTE2 maintains a current estimate $\pi^{(k)} \in \Pi$, and updates $\pi^{(k)}$ following three steps.

##### Step 1

The first step is to solve the linear program

$$
\begin{aligned}
\tilde{\pi}^{(k)} = \min_{\pi} \quad & \langle \nabla F(\pi^{(k)}), \pi \rangle_F \\
s.t. \pi \geq \mathbf{0} \\
& \pi \mathbf{1}_{n'} \leq g \\
& \pi^T \mathbf{1}_n \leq g' \\
& \mathbf{1}_n^T \pi \mathbf{1}_{n'} = s
\end{aligned}
$$

where gradient $\nabla F(\pi^{(k)})$ of $F(\pi^{(k)})$ is

$$
\nabla F(\pi^{(k)}) = (1-\alpha)\mathbf{C} + 2\alpha \mathbf{L}(D, D') \otimes \pi^{(k)} \tag{6}
$$

Notice that $\nabla F(\pi^{(k)})$ is a constant matrix with respect to $\pi$, and thus the linear program above is an instance of the partial Wasserstein optimal transport problem [9]. We follow [11] to compute the partial Wasserstein transport plan. Specifically, we transform the partial problem into a standard, full Wasserstein problem by adding a virtual spot to each of $g$ and $g'$ and modify the transport cost matrix $\nabla F(\pi^{(k)})$ accordingly such that the partial transport plan can be extracted from the extended transport matrix by removing the last column and last row. More details can be found in [11]. We solve the extended standard Wasserstein problem using the algorithm proposed in [6] as implemented in the Python Optimal Trasnport library [18].

**Step 2**

The second step finds the step size to move along the descent direction $\tilde{\pi}^{(k)}$ found in Step 1. That is, we find a $\gamma^{(k)}$ satisfying

$$
\gamma^{(k)} = argmin_{\gamma \in [0,1]} F(\pi^{(k)} + \gamma(\tilde{\pi}^{(k)} - \pi^{(k)})) \tag{7}
$$

Define $E^{(k)} = \tilde{\pi}^{(k)} - \pi^{(k)}$ and a function $\Phi : [0,1] \to \mathbb{R}$ such that

$$
\Phi(\gamma) = F(\pi^{(k)} + \gamma(\tilde{\pi}^{(k)} - \pi^{(k)})) \tag{8}
$$

We want to minimize $\Phi(\gamma)$ on $[0,1]$. We can rewrite $\Phi(\gamma)$ as

$$
\begin{aligned}
\Phi(\gamma) &= F(\pi^{(k)} + \gamma(\tilde{\pi}^{(k)} - \pi^{(k)})) \\
&= F(\pi^{(k)} + \gamma E^{(k)}) \\
&= (1-\alpha)\langle \mathbf{C}, \pi^{(k)} + \gamma E^{(k)} \rangle_F + \alpha \langle \mathbf{L}(D, D') \otimes (\pi^{(k)} + \gamma E^{(k)}), \pi^{(k)} + \gamma E^{(k)} \rangle_F \\
&= (1-\alpha)(\langle \mathbf{C}, \pi^{(k)} \rangle_F + \gamma \langle \mathbf{C}, E^{(k)} \rangle_F) + \alpha(\gamma^2 \langle \mathbf{L}(D, D') \otimes E^{(k)}, E^{(k)} \rangle_F \\
&\quad + 2\gamma \langle \mathbf{L}(D, D') \otimes E^{(k)}, \pi^{(k)} \rangle_F + \langle \mathbf{L}(D, D') \otimes \pi^{(k)}, \pi^{(k)} \rangle_F) \\
&= (1-\alpha)\langle \mathbf{C}, \pi^{(k)} \rangle_F + \gamma(1-\alpha)\langle \mathbf{C}, E^{(k)} \rangle_F + \gamma^2 \alpha \langle \mathbf{L}(D, D') \otimes E^{(k)}, E^{(k)} \rangle_F \\
&\quad + \gamma 2\alpha \langle \mathbf{L}(D, D') \otimes E^{(k)}, \pi^{(k)} \rangle_F + \alpha \langle \mathbf{L}(D, D') \otimes \pi^{(k)}, \pi^{(k)} \rangle_F \\
&= \gamma^2 \alpha \langle \mathbf{L}(D, D') \otimes E^{(k)}, E^{(k)} \rangle_F + \gamma((1-\alpha)\langle \mathbf{C}, E^{(k)} \rangle_F + 2\alpha \langle \mathbf{L}(D, D') \otimes E^{(k)}, \pi^{(k)} \rangle_F) \\
&\quad + (1-\alpha)\langle \mathbf{C}, \pi^{(k)} \rangle_F + \alpha \langle \mathbf{L}(D, D') \otimes \pi^{(k)}, \pi^{(k)} \rangle_F \\
&= a\gamma^2 + b\gamma + c
\end{aligned}
$$

where $a, b, c$ are constants calculated from known quantities

$$
\begin{aligned}
a &= \alpha\langle \mathbf{L}(D, D') \otimes E^{(k)}, E^{(k)}\rangle_F \\
b &= (1-\alpha)\langle \mathbf{C}, E^{(k)}\rangle_F + 2\alpha\langle \mathbf{L}(D, D') \otimes E^{(k)}, \pi^{(k)}\rangle_F \\
c &= (1-\alpha)\langle \mathbf{C}, \pi^{(k)}\rangle_F + \alpha\langle \mathbf{L}(D, D') \otimes \pi^{(k)}, \pi^{(k)}\rangle_F
\end{aligned}
$$

Now, minimizing $\Phi(\gamma)$ on $[0, 1]$ is just minimizing a univariate quadratic function on $[0, 1]$, which can be done by testing the convexity and finding the axis of symmetry.

**Step 3**

$\tilde{\pi}^{(k)}$ is calculated in step 1. $\gamma^{(k)}$ is calculated in step 2. Now update

$$\pi^{(k+1)} = \pi^{(k)} + \gamma^{(k)}(\tilde{\pi}^{(k)} - \pi^{(k)}) \tag{9}$$

In practice, we test convergence by comparing the difference between the objective cost of $\pi^{(k)}$ and $\pi^{(k+1)}$ to a small constant. Algorithm 1 shows the pseudocode of our conditional gradient algorithm to optimize the partial-FGW objective.

**Algorithm 1: Conditional gradient algorithm for partial-FGW**

**Input:** Transport cost matrix $\mathbf{C}$; pairwise cost tensor $\mathbf{L}(D, D')$; feasible region $\mathcal{P}$; balance parameter $\alpha$; convergence parameter $\delta$

**Initialize** initial guess $\pi^{(0)} \in \Pi$;

**while** $F(\pi^{(k+1)}) - F(\pi^{(k)}) > \delta$ **do**

- $\nabla F(\pi^{(k)}) = (1-\alpha)\mathbf{C} + 2\alpha \mathbf{L}(D, D') \otimes \pi^{(k)}$ `// Gradient computation`
- $\tilde{\pi}^{(k)} = argmin_{\pi \in \Pi} \langle \nabla F(\pi^{(k)}), \pi\rangle_F$ `// Step 1: Solve partial-W subproblem`
- $\gamma^{(k)} = argmin_{\gamma \in [0,1]} F(\pi^{(k)} + \gamma(\tilde{\pi}^{(k)} - \pi^{(k)}))$ `// Step 2: Line search`
- $\pi^{(k+1)} = \pi^{(k)} + \gamma^{(k)}(\tilde{\pi}^{(k)} - \pi^{(k)})$ `// Step 3: Update`

**return** $\pi^{(k)}$

#### 4 The histological image dissimilarity matrix

The H&E image associated with each slice can be represented by a matrix $H \in \mathbb{N}^{n \times 3}$, where $n$ is the number of spots on the slice and the $i$-th row $\mathbf{h}_{i\cdot}$ is the RGB value of the pixel of spot $i$ in the H&E image. In reality, a spot may occupy a circle instead of a pixel in the image, so we take the average value of all pixels in the circle as the RGB value for the spot. Given two ST slices $(X, D, H)$ and $(X', D', H')$, we integrate the H&E image information into the partial-FGW framework by defining a cost matrix $\mathbf{C}_{image} \in \mathbb{R}^{n \times n'}$ to encode the Euclidean distance between the RGB value of each spot of the first slice and the RGB value of each spot of the second slice, and spots with similar histology achieve lower costs. That is, $[\mathbf{C}_{image}]_{ij} = \|\mathbf{h}_{i\cdot} - \mathbf{h}'_{j\cdot}\|_2$. If PASTE2 aligns spot $i$ to spot $j$, then both the gene expression profiles and the histology RGB values of spot $i$ and $j$ should be similar.

#### 5 Optimal projection and 3D reconstruction

Given a series of consecutive slices $(X^{(1)}, Z^{(1)}), ..., (X^{(t)}, Z^{(t)})$, where $X$ is the gene expression matrix and $Z$ is the 2D location matrix, for $k = 1, ..., t-1$, we seek to project the coordinates of slice $k+1$ onto the coordinates of slice $k$ such that the partial alignment $\pi^{(k)}$ between the two slices is respected. The projection is defined by a rotation matrix $R \in \mathbb{R}^{2\times 2}$ and a translation vector $t \in \mathbb{R}^2$ that is applied to the spatial coordinates $Z^{(k+1)}$ of slice $k+1$. The derivation here is similar to the 3D reconstruction in PASTE, but can handle partial alignment matrices.

Given ST slices with spatial coordinates $Z \in \mathbb{R}^{2\times n}$ and $W \in \mathbb{R}^{2\times n'}$, and a partial alignment $\pi \in \mathcal{P}(g, g', s)$ between the two slices, we want to find a rotation matrix $R \in \mathbb{R}^{2\times 2}$ and a translation vector $t \in \mathbb{R}^2$ for $W$ that minimizes

$$Q(t, R) = \sum_{i,j} \pi_{ij} \|Z_{\cdot i} - RW_{\cdot j} - t\|^2 \tag{10}$$

We first show that we can assume no translation is needed ($t = 0$) by scaling both $Z$ and $W$. Assume $R$ is fixed, we take the derivative of $Q$ with respect to $t$ and compare to 0

$$\begin{aligned}
\frac{\partial Q}{\partial t} &= -2\sum_{i,j} \pi_{ij}(Z_{\cdot i} - RW_{\cdot j} - t) \\
&= -2\sum_{i} Z_{\cdot i}\sum_{j} \pi_{ij} + 2\sum_{j} RW_{\cdot j}\sum_{i} \pi_{ij} + 2t\sum_{i,j} \pi_{ij} \\
&= -2\sum_{i} Z_{\cdot i}p_i + 2\sum_{j} RW_{\cdot j}q_j + 2ts \\
&= -2\sum_{i} Z_{\cdot i}p_i + 2R\sum_{j} W_{\cdot j}q_j + 2ts \\
&= -2Zp + 2RWq + 2ts = 0
\end{aligned}$$

where $p = \pi\mathbf{1}_{n'}, q = \pi^T\mathbf{1}_n, s = \mathbf{1}_n^T\pi\mathbf{1}_{n'}$. We have $t = \frac{1}{s}(Zp - RWq)$. Then, substitute $t = \frac{1}{s}(Zp - RWq)$ into $Q$, we get

$$\begin{aligned}
Q(t, R) &= \sum_{i,j} \pi_{ij}\|Z_{\cdot i} - RW_{\cdot j} - t\|^2 \\
&= \sum_{i,j} \pi_{ij}\|Z_{\cdot i} - RW_{\cdot j} - \frac{1}{s}Zp + \frac{1}{s}RWq\|^2 \\
&= \sum_{i,j} \pi_{ij}\|(Z_{\cdot i} - \frac{1}{s}Zp) - R(W_{\cdot j} - \frac{1}{s}Wq)\|^2
\end{aligned}$$

Since $\frac{1}{s}Zp$ and $\frac{1}{s}Wq$ does not depend on $R$, if we replace $Z_{\cdot i}$ with $Z_{\cdot i} - \frac{1}{s}Zp$ and $W_{\cdot j}$ with $W_{\cdot j} - \frac{1}{s}Wq$, $Q$ is minimized with respect to $t$ and we only need to find the optimal rotation $R$. Hence we can assume no translation is needed by scaling both $R$ and $W$.

To find the optimal rotation $R$, rewrite $Q$ in matrix notation

$$
\begin{aligned}
Q(R) &= \sum_{i,j} \pi_{ij} \|Z_{\cdot i} - RW_{\cdot j} - t\|^2 \\
&= \sum_{i,j} \pi_{ij} (Z_{\cdot i} - RW_{\cdot j} - t)^T (Z_{\cdot i} - RW_{\cdot j} - t) \\
&= \sum_{i,j} \pi_{ij} (Z_{\cdot i}^T Z_{\cdot i} - Z_{\cdot i}^T RW_{\cdot j} - W_{\cdot j}^T R Z_{\cdot i} + W_{\cdot j}^T R^T R W_{\cdot j}) \\
&= -2 \sum_{i,j} \pi_{ij} (Z_{\cdot i}^T RW_{\cdot j}) + \alpha \\
&= -2 \operatorname{Tr}(Z^T RW\pi^T) + \alpha \\
&= -2 \operatorname{Tr}(RW\pi^T Z^T) + \alpha
\end{aligned}
$$

where $\alpha$ is a constant independent of $R$. Let $U\Sigma V^T$ be the SVD decomposition of the matrix $W\pi^T Z^T$. Then,

$$
\begin{aligned}
Q(R) &= -2 \operatorname{Tr}(RW\pi^T Z^T) + \alpha \\
&= -2 \operatorname{Tr}(RU\Sigma V^T) + \alpha \\
&= -2 \operatorname{Tr}(\Sigma V^T RU) + \alpha
\end{aligned}
$$

Since $V, R, U$ are all orthonormal matrices, and $\Sigma$ is a diagonal matrix with positive entries, the maximum of $\operatorname{Tr}(\Sigma V^T RU)$, hence the minimum of $Q$, will be achieved when $V^T RU = I$. Therefore, the optimal rotation that minimizes $Q$ is $R = VU^T$.

By finding the optimal projection for each slice pair, we project each slice onto the same 2D coordinate grid and create a common coordinate system for all slices in 3D where the z-axis is determined by the actual distance between each slice in the tissue.

#### 6 Simulation procedure

The simulated partial slices are based on DLPFC sample 151674, where each spot is labeled with a manual cortical layer annotation from [32]. This slice contains 3635 spots and 12381 genes after filtering out all the spots and genes with less than 100 transcript counts overall. We generated two partially overlapping subslices from this slice in the following way.

1. Let $s$ be a percentage number between 0 and 100 (or equivalently, fraction number between 0 and 1.) We choose two horizontal lines on the slice, line 1 and line 2, with line 1 below line 2, that generates two subslices. The upper subslice is the subslice above line 1, and the lower subslice is the subslice below line 2. The upper and lower subslice overlap at $s$ percent of their spots. $s$ controls the exact locations of line 1 and line 2 that cuts the slice into subslices.

2. For each spot $i$ in the lower subslice, we resample its gene expression profile as follows. Let $v_i$ be the original gene expression vector of spot $i$, and $\mu_i = \sum_i v_i$ be the total read count. Let $\delta$ be a small pesudocount. We resample $v_i$ according to a multinomial distribution $v_i \sim Multinomial(\mu_i, \frac{v_i + (\delta \cdot 0.0002 \cdot \mu_i) \cdot \mathbf{1}_p}{\mu_i + (\delta \cdot 0.0002 \cdot \mu_i) \cdot p})$, where $p$ is the number of genes and $\mathbf{1}_p$ is an all-one vector of length $p$.

That is, for each spot in the bottom slice, we add a small pseudocount to each gene in proportion to the total read counts of the spot, and then resample the expression profile under a multinomial distribution defined by the read counts of each gene after the addition of the pseudocount. This way, we have a different pseudocount for each spot proportional to the spot's total read count so that the noises we introduce are the same across spots. We choose the ratio as $0.0002 \cdot \mu_i$ for a spot with read count $\mu_i$ so that the median pseudocount added across spots would be $\delta$ because the median read counts for each spot in this slice is about 5000.

### 7 Label Transfer Adjusted Rand Index (LTARI)

We evaluate the alignment accuracy for partial slice alignments using what we call the Label Transfer Adjusted Rand Index (LTARI). LTARI is a score that measures the ability of the alignment to transfer labels of the aligned region from one slice to the other. Intuitively, a good partial alignment would find and align spots with the same cell type, hence we define a score that measures the alignment accuracy as the agreement of cell type labels of aligned pairs of spots. Note that the ground truth annotation of cell type of each spot should be available for calculating LTARI. Specifically, for each spot $j$ in the second slice that is aligned by PASTE2, the alignment induces a new cell type label for the spot by assigning it $\ell(j) = \ell(\mathrm{argmax}_i\, \pi_{ij})$, the label of the spot $i$ in the first slice that achieves the highest $\pi_{ij}$ over all the spots in the first slice. That is, we assign each aligned spot in the second slice the label of the spot in the first slice that is mostly likely aligned to it according to the computed alignment. This assignment transfers the labels of spots from the first slice to the second slice in the aligned region. We then compare this transferred labeling with the ground truth labeling of the aligned region of the second slice and compute the ARI of the two clusterings. For PASTE2, the Label Transfer ARI is calculated on the region that PASTE2 chooses to align since not all spots receive an alignment, while for PASTE the Label Transfer ARI is calculated for the entire slice because PASTE have to align every spot to some spot in the other slice. A high LTARI indicates that the (partial) alignment tends to align each spot to some spot on the other slice with the same cell type label, hence corresponds to a better alignment. Notice that LTARI can be defined in the opposite direction, comparing the transferred labeling with the ground truth labeling of the first slice, but in practice we do not observe a significant gap between the LTARI of the two directions.

### 8 Benchmarking PASTE2 with other methods

We benchmarked PASTE2 against PASTE, Pamona, and Tangram on the DLPFC dataset. Both PASTE2 and Pamona are partial alignment methods that can handle partially overlapped datasets, while PASTE and Tangram assume the two datasets to align have the same underlying cellular structure. We treated both ST slices as scRNA-seq datasets for Pamona by dropping the information about spatial coordinates. Since Pamona is a partial alignment algorithm, it takes as input the number of shared cells between the two datasets, and we provided the ground-truth number of spots (70% of total spots) in the overlap region of the two slices. To run Tangram, we treated the first slice as a scRNA-seq dataset and mapped its spots onto the second ST slice. We used the uniform density prior for Tangram such that the mapping returned by Tangram will have uniform marginals over each spot as in PASTE. We ran Tangram for 500 iterations instead of the default 1000 because empirically the loss does not change much after 500 iterations. We ran PASTE2 with the ground-truth overlap percentage $s$ as well.

### 9 Evaluation of the estimation of overlap percentages

We evaluated PASTE2's model selection procedure (Supplement §1) to estimate the overlap percentage $s$ on the simulated dataset described in §3.1 and the real ST slices described in §3.2. For the simulated dataset, we used PASTE2 to estimate the overlap percentage of simulated pairs of slices which overlap at 90%, 70%, 50%, and 30% of their areas, and with pseudocount added to the gene expression data $\delta = 0.1, 1.0, 2.0, 3.0$. For the real slices, we used PASTE2 to estimate the overlap percentage of all pairs of adjacent subslices of all individuals analyzed in §3.2, with each pair roughly overlap at 70% of their areas.

The estimation result on simulated pairs of slices indicates that when the overlap between the two slices is high ($> 50\%$), PASTE2 estimates the overlap percentage accurately, with all estimations within 10% of the ground truth even when the noise level $\delta = 3.0$ (Fig. S12a). When the overlap is less than 50%, PASTE2 still correctly recovers the overlap when $\delta$ is reasonably small.

On pairs of DLPFC subslices, both horizontally and vertically overlapping, for 8 out of 18 pairs, the PASTE2 estimation of overlap percentage is within 10% of the reference overlap, and the estimations of 16 out of 18 pairs are within 20% of the reference overlap (Fig. S12b). However, we note that the 70% reference overlap simply means that the rectangular boxes used to crop out subslices, as shown in Fig. S8, have about 70% of their areas overlapping between each pair. Due to factors such as variations in shapes of the tissue and different geometries of each layer in different slices, the true overlap percentage may differ from 70%. For example, comparing the 3D reconstruction (optimal projection) of pair BC of sample 1 based on an alignment with 70% of overlap and the 3D reconstruction of the same pair based on an alignment with 30% of overlap (estimated by PASTE2), it is clear that the PASTE2 estimation of 30% leads to a better reconstruction than the reference 70% (Fig. S13), and the alignment LTARI increases from 0.07 to 0.18. Additionally, we want to mention that the differences in geometries of layers across slices, complicated with technical artifacts such as sharp difference in UMI counts across layers, may result in model unindentifiability issues.

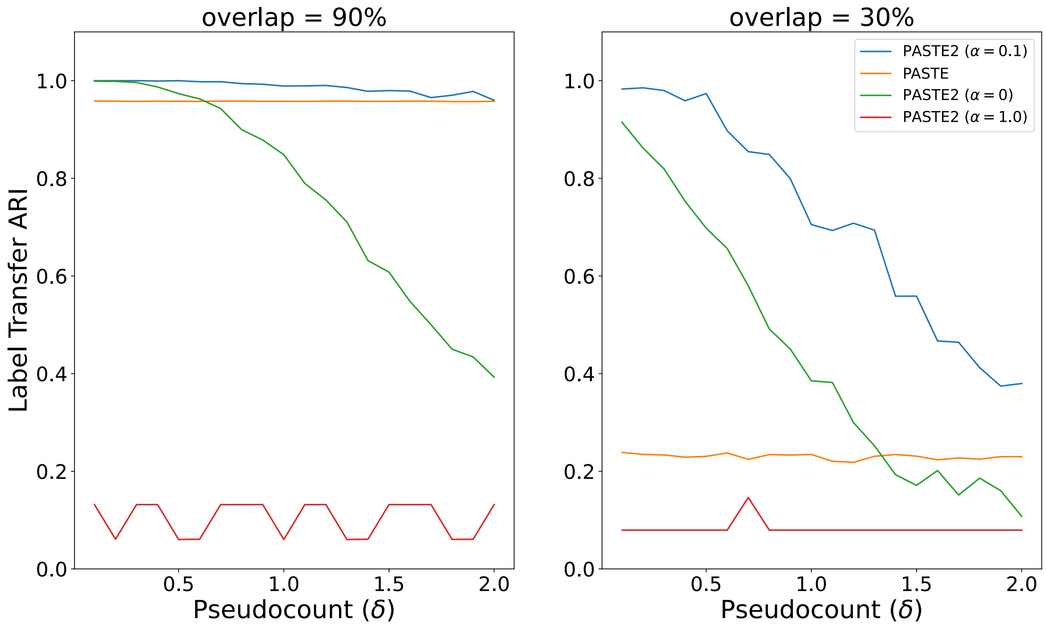


**Figure 6: Additional PASTE2 results on simulated data.** Label Transfer ARI of the alignment result versus the added pseudocount ($\delta$) for PASTE2 $\alpha = 0$ (gene expression information only), PASTE2 $\alpha = 1$ (spatial information only), PASTE2 $\alpha = 0.1$ (both), and PASTE (full alignment), for overlap percentages 30% and 90%.

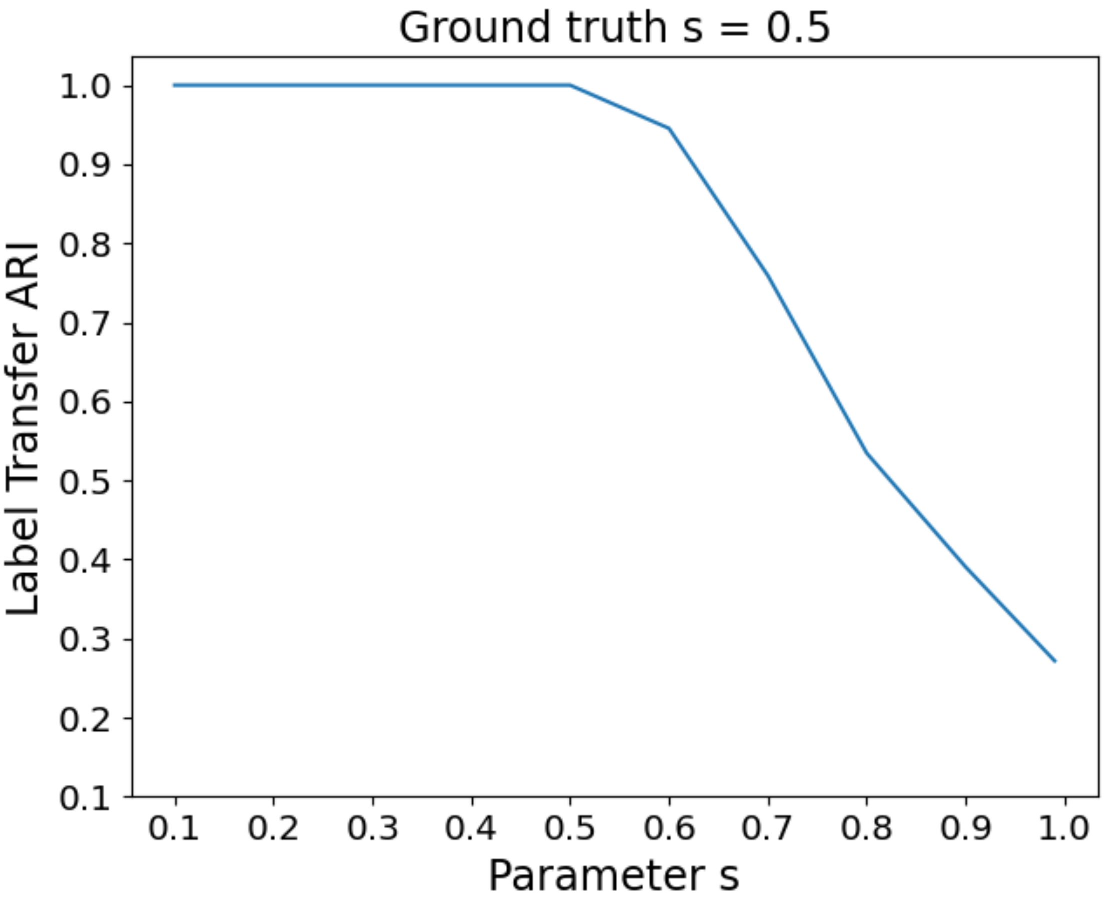


**Figure 7: Effect of the overlap percentage parameter $s$ on PASTE2 results.** Label Transfer ARI of the PASTE2 alignment result for varying values of the parameter $s$ on a simulated pair with overlap percentage $s = 0.5$.

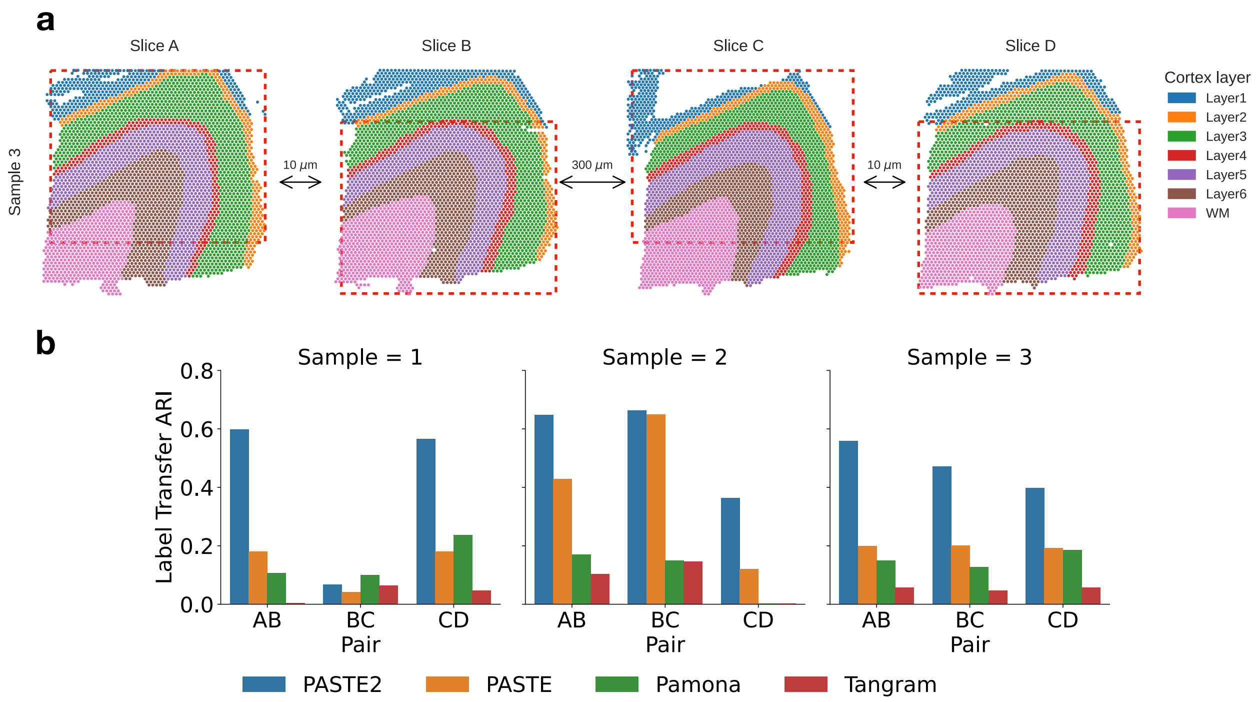


**Figure 8: Horizontal partial alignment tasks.** **a**, Horizontal (red bounding boxes) subslices cropped out of four slices from sample 3. Each pair of adjacent subslices overlap at 70% of the areas. **b**, LTARI of alignments of each pair of adjacent horizontal subslices for PASTE2, PASTE, Pamona, and Tangram.

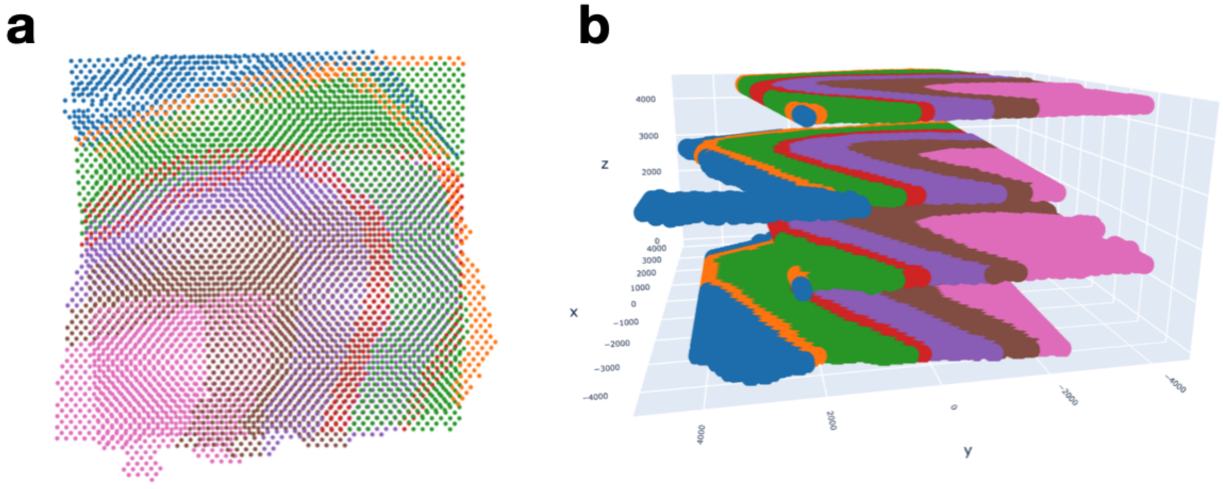


**Figure 9: Spatial reconstruction of DLPFC slices.** **a**, Optimal projection of vertical subslices of sample 3 slice AB based on PASTE alignment. **b**, PASTE2 3D reconstruction of the tissue of sample 3 from four horizontal partial slices. Note that $z$-axis is not to scale.

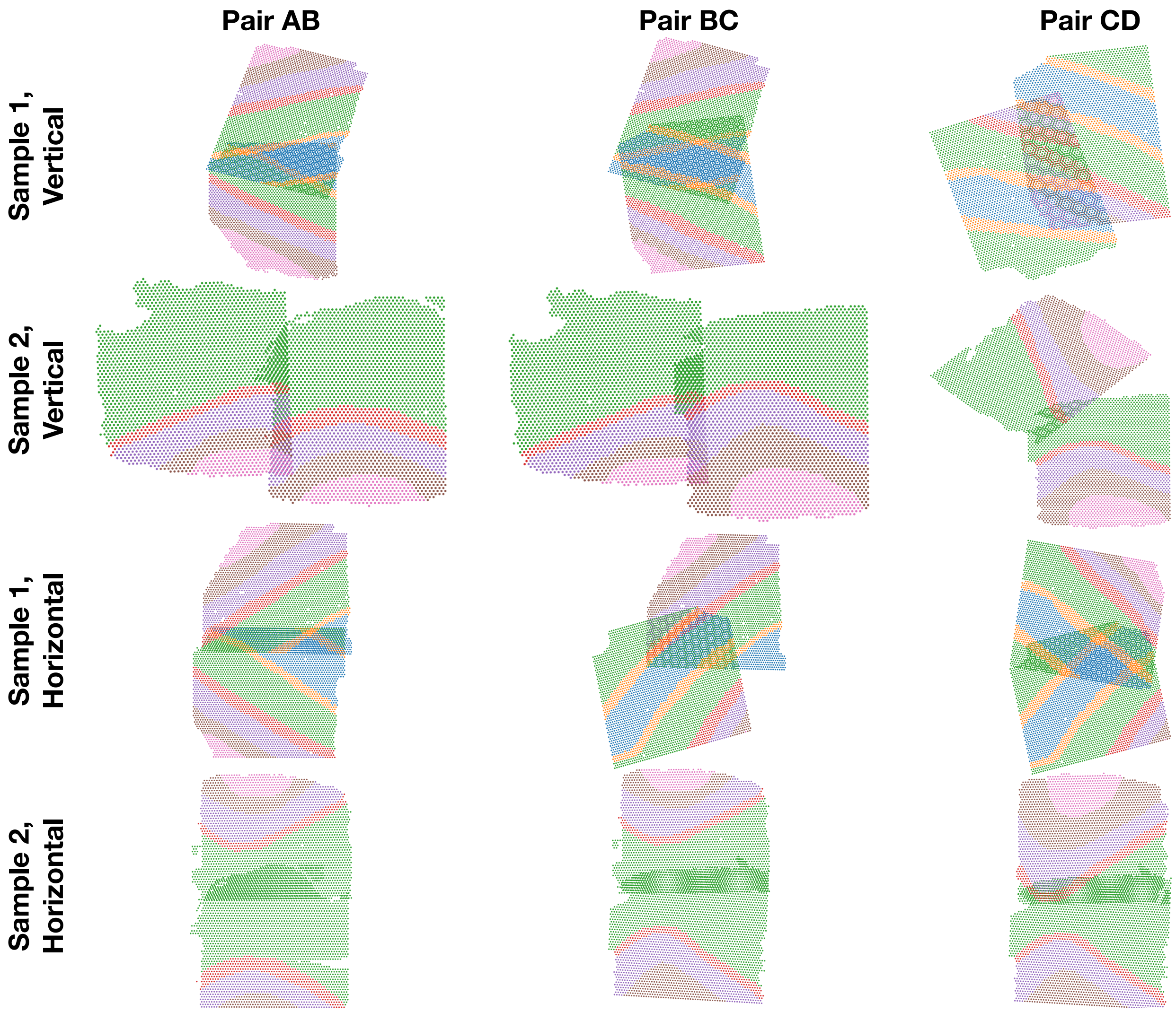


**Figure 10: Alignment results of STUtility on partial DLPFC slices.** Each pair of adjacent vertical and horizontal partial slices aligned by STUtility. Each pair is plotted according to the aligned coordinates output by STUtility.

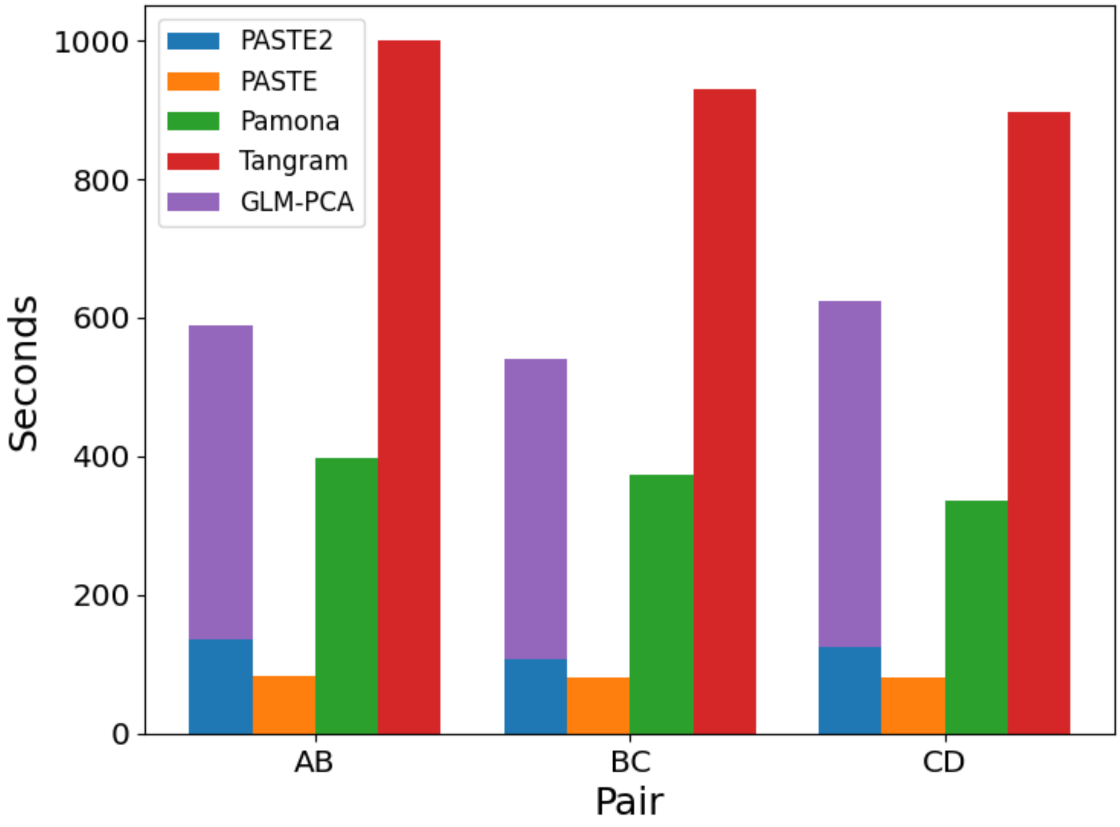


**Figure 11: Running time of four methods on DLPFC sample 3.** The running time of PASTE2, PASTE, Pamona, and Tangram on the vertical subslice pairs of DLPFC sample 3. The running time of PASTE2 is broken into two parts: time spent on the GLM-PCA subroutine by calling another library, and the time spent on the PASTE2 conditional gradient optimization procedure.

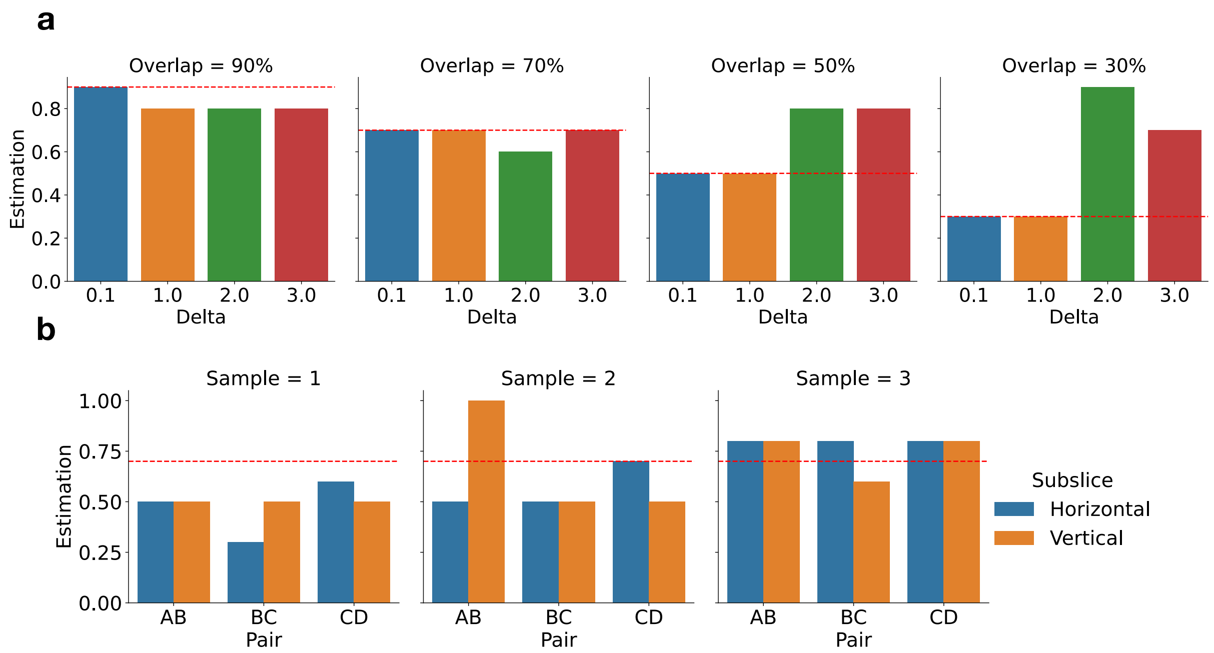


**Figure 12: PASTE2 estimation of overlap percentages on simulated and real DLPFC slices. a**, PASTE2 estimation of overlap percentages for simulated pairs of slices with four different overlap percentages, each with four gene expression noise levels. The red dotted line denotes the ground truth. **b**, PASTE2 estimation of overlap percentages of horizontal and vertical subslices cropped from real DLPFC slices. The red dotted line denotes the 70% reference overlap.

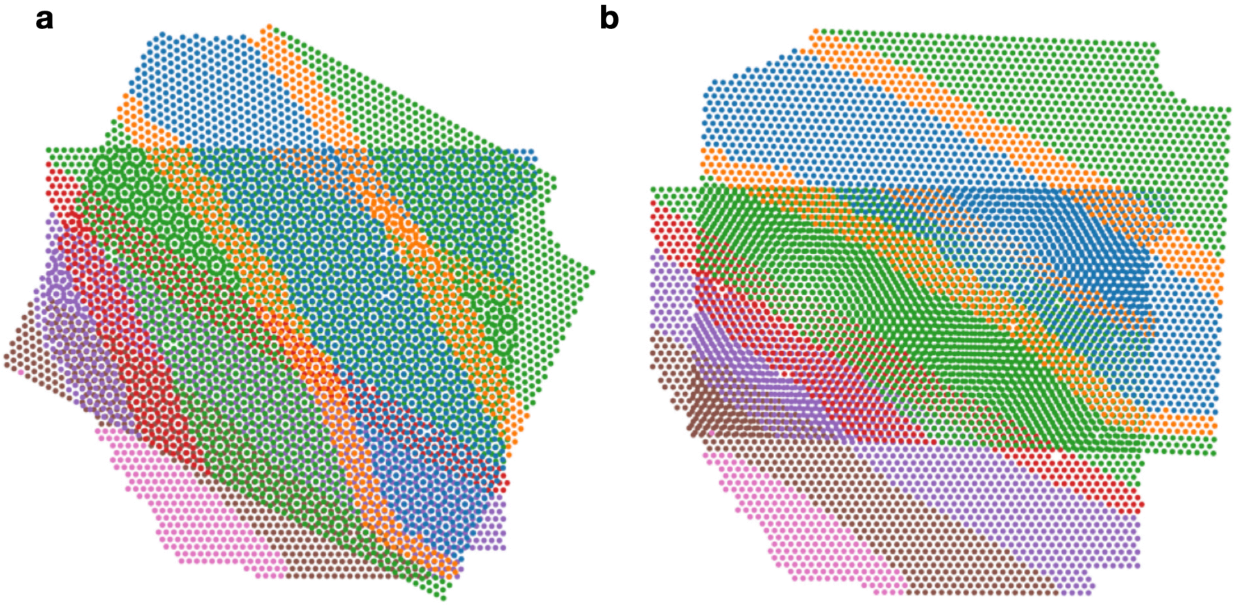


**Figure 13: PASTE2 3D reconstruction of sample 1 pair BC. a**, Reconstruction based on PASTE2 alignment with input $s = 0.7$. **b**, Reconstruction based on PASTE2 alignment with input $s = 0.3$.

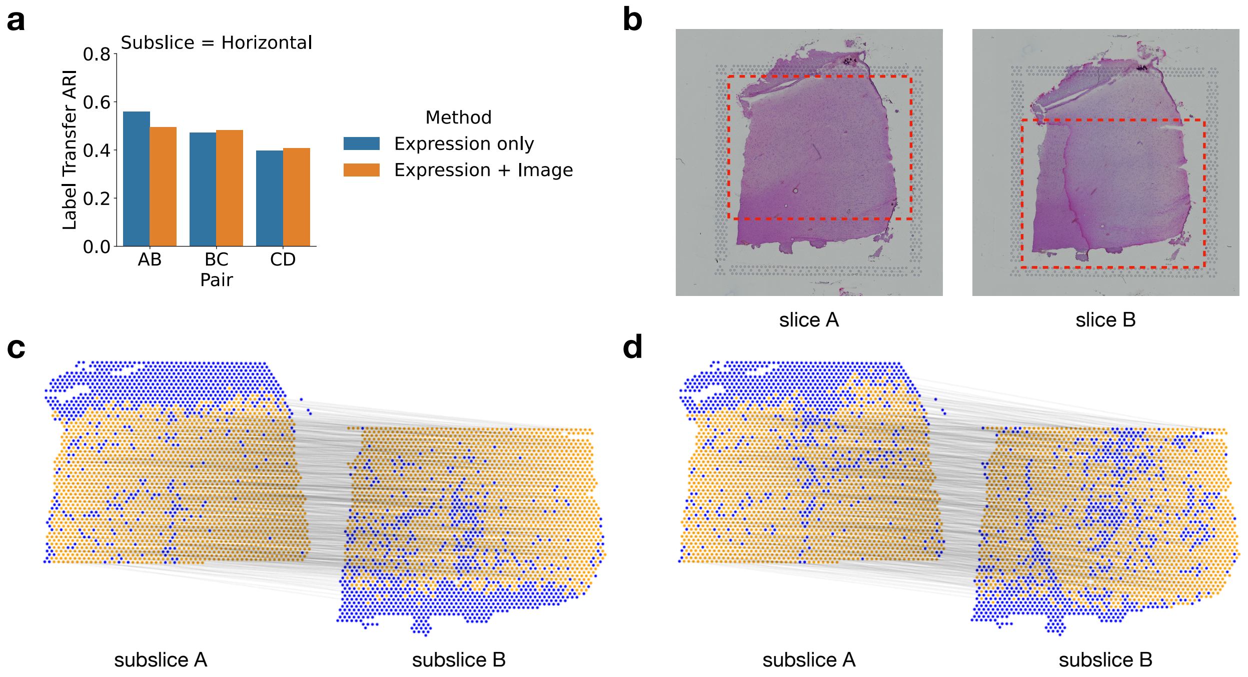


**Figure 14: Effect of histological image information on alignment of horizontal subslices of sample 3. a**, Comparing LTARI of PASTE2 alignments using expression information only with both expression and image information, for horizontal subslices of DLPFC sample 3. Spatial information is used in both modes. **b**, Histological images of sample 3 slice A and slice B. The red boxes bound the horizontal subslices cropped for partial alignment. The lower part of subslice A overlaps with the upper part of slice B. **c**, Visualization of PASTE2 alignment of the subslice pair when using gene expression information only. Yellow spots are spots that PASTE2 chooses to align, and blue spots are decided non-overlapping. The black lines connect pairs of spots aligned by PASTE2 with high weight. **d**, Visualization of PASTE2 alignment of the subslice pair when using both gene expression and histological image information.

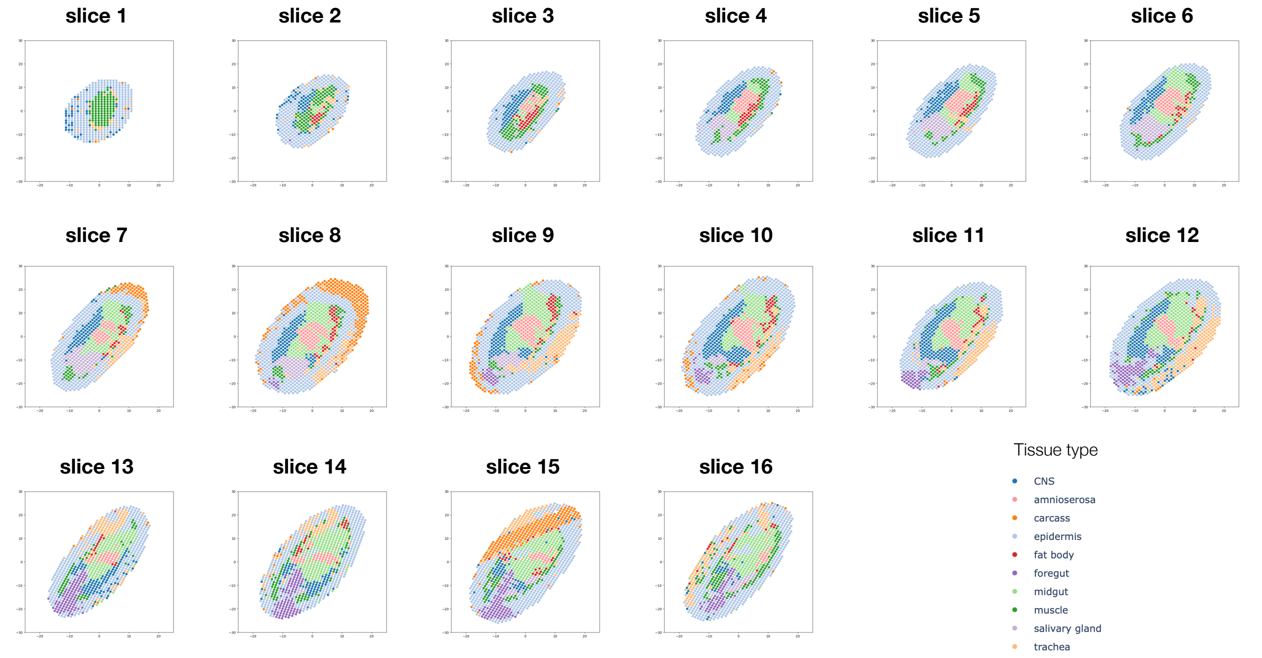


**Figure 15: Stereo-seq slices of E14-16 Drosophila embryo.** Visualization of 16 slices of E14-16 Drosophila embryo. Coloring of spots is according to cell type annotations in [47].

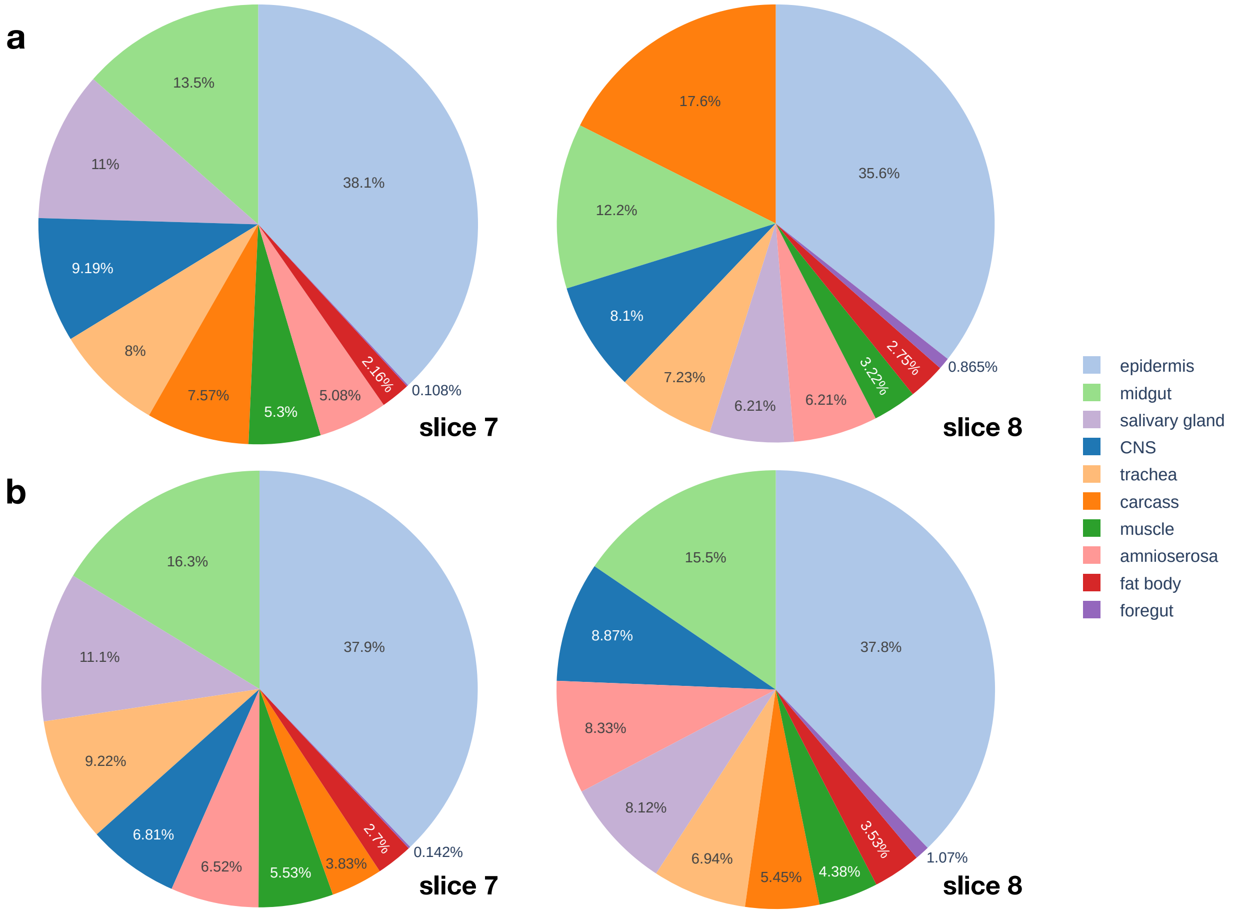


**Figure 16: Cell type compositions of slice 7 and slice 8 before and after alignment.** **a**, Cell type compositions of original slice 7 and slice 8. Both carcass and salivary gland cells show large imbalance between the two slices. **b**, Cell type compositions of the aligned regions in slice 7 and slice 8.

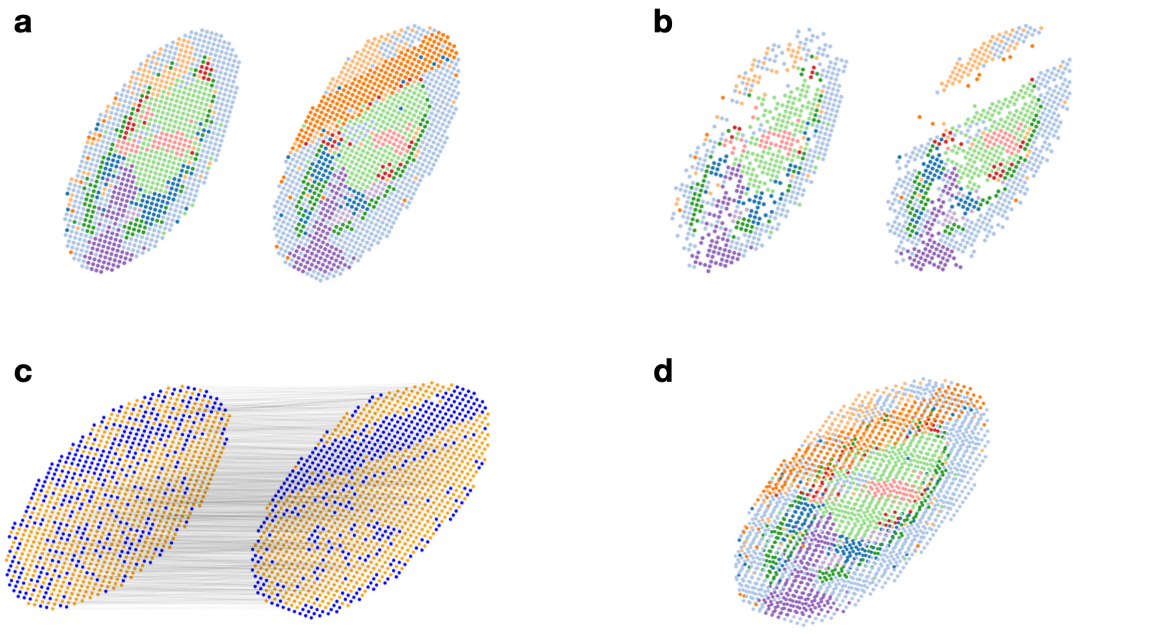


**Figure 17: PASTE2 alignment of Drosophila embryo slice 14 and slice 15.** **a**, Visualization of the two slices before alignment. Slice 15 has a stripe of carcass cells (orange color) that is not present in slice 14. **b**, Visualization of the spots from the two slices that are chosen to be aligned by PASTE2. The orange spots are left out. **c**, The PASTE2 alignment. Yellow spots are spots that PASTE2 chooses to align, and blue spots are decided non-overlapping. The black lines denote the actual spot-spot alignment. **d**, Optimal projection of slice 14 onto slice 15 based on the PASTE2 alignment.

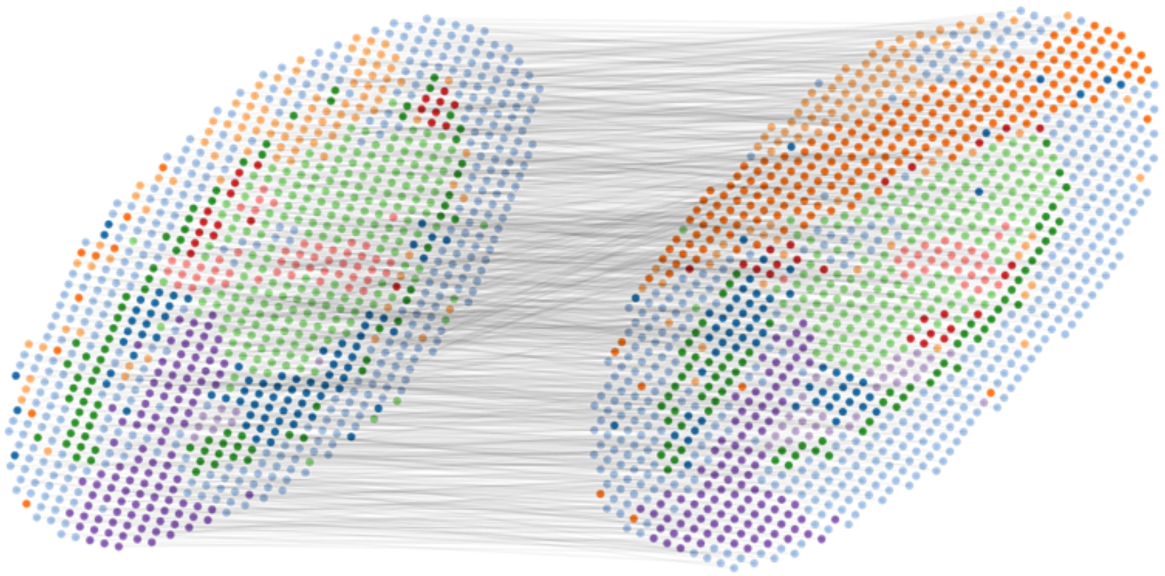

**Figure 18: PASTE alignment of Drosophila embryo slice 14 and slice 15.** The black lines denote the actual spot-spot alignment. The carcass (orange) spots on slice 15 are mapped arbitrarily.

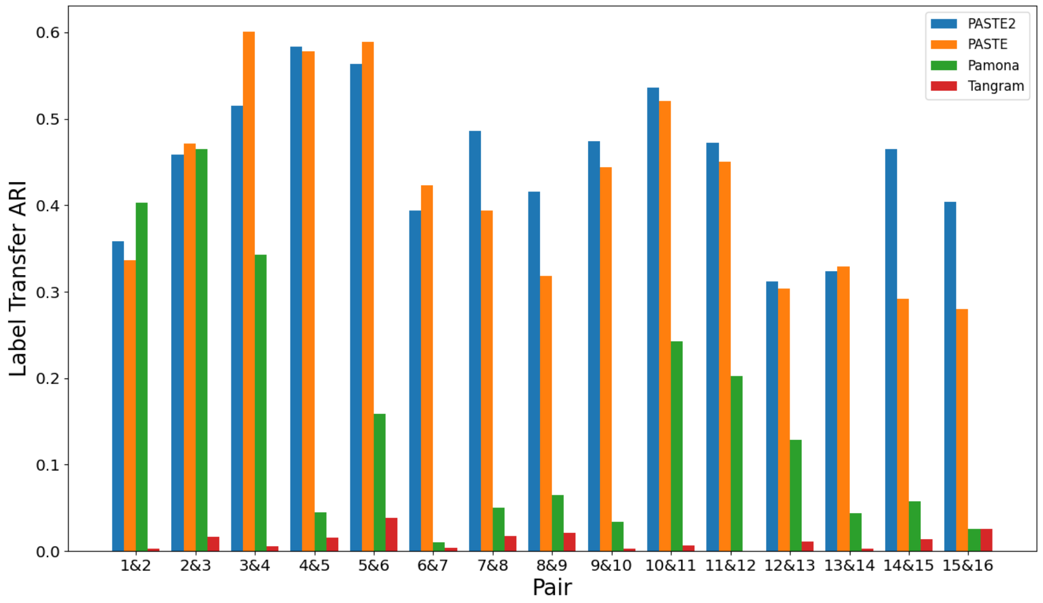


**Figure 19: Comparison of the results of four methods aligning each pair of adjacent slices of the Drosophila embryo.** LTARI of pairwise alignments computed by PASTE2, PASTE, Pamona, and Tangram for each pair of adjacent slices.
